# Single-Cell Protein Atlas of Transcription Factors Reveals the Combinatorial Code for Spatiotemporal Patterning the *C. elegans* Embryo

**DOI:** 10.1101/2020.06.30.178640

**Authors:** Xuehua Ma, Zhiguang Zhao, Long Xiao, Weina Xu, Yangyang Wang, Yanping Zhang, Gang Wu, Zhuo Du

## Abstract

A high-resolution protein atlas is essential for understanding the molecular basis of biological processes. Using protein-fusion reporters and imaging-based single-cell analyses, we present a protein expression atlas of *C. elegans* embryogenesis encompassing 266 transcription factors (TFs) in nearly all (90%) lineage-resolved cells. Single-cell analysis reveals a combinatorial code and cascade that elucidate the regulatory hierarchy between a large number of lineage-, tissue-, and time-specific TFs in spatiotemporal fate patterning. Guided by expression, we identify essential functions of CEH-43/DLX, a lineage-specific TF, and ELT-1/GATA3, a well-known skin fate specifier, in neuronal specification; and M03D4.4 as a pan-muscle TF in converging muscle differentiation in the body wall and pharynx. Finally, systems-level analysis of TF regulatory state uncovers lineage- and time-specific kinetics of fate progression and widespread detours of the trajectories of cell differentiation. Collectively, our work reveals a single-cell molecular atlas and general principles underlying the spatiotemporal patterning of a metazoan embryo.

## INTRODUCTION

How a totipotent zygote generates all specialized cell types is a central question in developmental biology. *Caenorhabditis elegans* (*C. elegans*) is the first multicellular organism to have its cell atlas completely mapped, including the lineage history, anatomy, and fate of all 959 somatic cells (Albertson and Thomson, 1976 ; Hall and Altun, 2008; Sulston and Horvitz, 1977; Sulston et al., 1983; White et al., 1986) and quantitative cellular behaviors during embryogenesis (Bao et al., 2006; Giurumescu et al., 2012; Moore et al., 2013; Richards et al., 2013; Schnabel et al., 2006), making it a leading model organism for elucidating the regulation of cell differentiation at single-cell level. However, whereas the cell atlas of development has been fully resolved, the molecular atlas of differentiation remains to be systematically elucidated. For example, we still lack comprehensive knowledge of the chromatin states, gene transcription, protein expression, molecular interactions, and regulatory networks within each developing cell, knowledge critical for the understanding, modeling, and engineering of cell fates.

Cell differentiation is driven by timely expression of specific gene sets in specific cells. Advances in single-cell genomics and live imaging-based single-cell analysis provide the opportunity to dissect gene expression programs in each cell during cell differentiation. Single-cell RNA-sequencing (scRNA-seq) allows transcriptome mapping of isolated single cells and inference of cell identity based on transcriptome similarity between cells and marker genes with known cell-specificities (Luecken and Theis, 2019). Using scRNA-seq, cellular transcriptional atlases have been mapped during early (Tintori et al., 2016) and throughout the whole of embryogenesis (Packer et al., 2019). While spectacular, the scRNA-seq approach has three limitations: temporal resolution is not high, cell lineage identities are only partially resolved (Kiselev et al., 2019), and mRNA expression correlates poorly with cellular protein abundance (Grun et al., 2014; Marguerat et al., 2012; Peshkin et al., 2015; Taniguchi et al., 2010).

Complementarily, 3D time-lapse imaging using fluorescent reporters combined with automated cell lineage tracing and fluorescence quantification allows precise measurement of the transcript or protein levels of specific genes in identity-resolved single cells at high temporal resolution (Bao et al., 2006; Du et al., 2014; Liu et al., 2009; Murray et al., 2008; Murray et al., 2012). Although only one gene is analyzed each time, this approach presents a unique advantage in tracing gene expression in lineaged cells at high temporal resolution (Murray et al., 2012), and has been used to map a cellular expression atlas of ∼100 regulatory genes. However, these pioneering studies have either focused on a static developmental stage (Liu et al., 2009) or cover only half of the embryo’s cells (Murray et al., 2012). Moreover, promoter-fusion rather than protein-fusion reporters were used to profile most mRNA levels.

Here, using an imaging-based approach, we present a dynamic single-cell protein atlas encompassing hundreds of transcription factors (TFs), those essential regulators of gene expression and cell differentiation. Using protein-fusion fluorescent reporters for 266 TFs (including 86 endogenous reporters generated here), we quantified and integrated TF protein expression in 1,204 lineaged embryonic cells, covering 90% of all cells generated during embryogenesis. Systematic single-cell analyses revealed a spatiotemporal combinatorial code and cascade of TFs that illuminate how lineage-, tissue-, and time-specific TFs drive hierarchical cell fate specification and differentiation. We further uncovered new functions of TFs in the lineage-and tissue-based regulation of cell fate and discovered general principles regarding the dynamics of cellular states during cell lineage differentiation. Our work contributes to a molecular and mechanistic understanding of cell differentiation at single-cell resolution.

## RESULTS

### Quantifying protein expression of TFs in nearly all lineage-traced embryonic cells

We selected 290 high-confidence target TFs, including 273 conserved in humans and 17 others of potential interest for *C. elegans* embryogenesis, which encompass diverse important TF classes such as the homeodomain, bHLH, forkhead, GATA, and ETS families (Figure 1A, Figure S1A, and Table S1). Protein-fusion GFP reporter strains were previously generated for 195 of these (Figure S1B), in which GFP was recombineered into the TF coding region in a fosmid vector and integrated into the genome at low copy number (Araya et al., 2014). Since the fosmids harbor 25-50 kb genomic regions (4-6 genes) including most regulatory elements, reporter expression highly recapitulates that of the endogenous protein (Sarov et al., 2012). For TFs not already having reporter strains, we performed CRISPR/Cas9- mediated gene knockin to tag endogenous genes with mNeonGreen/GFP, successfully constructing endogenous protein-fusion reporters for 86 TFs (Figure S1C). Together with previously generated knockin reporters for another eight TFs, we have collected 291 protein-fusion reporters for 266 TFs (Figure 1B, Figure S1D and Table S1).

**Figure 1.**
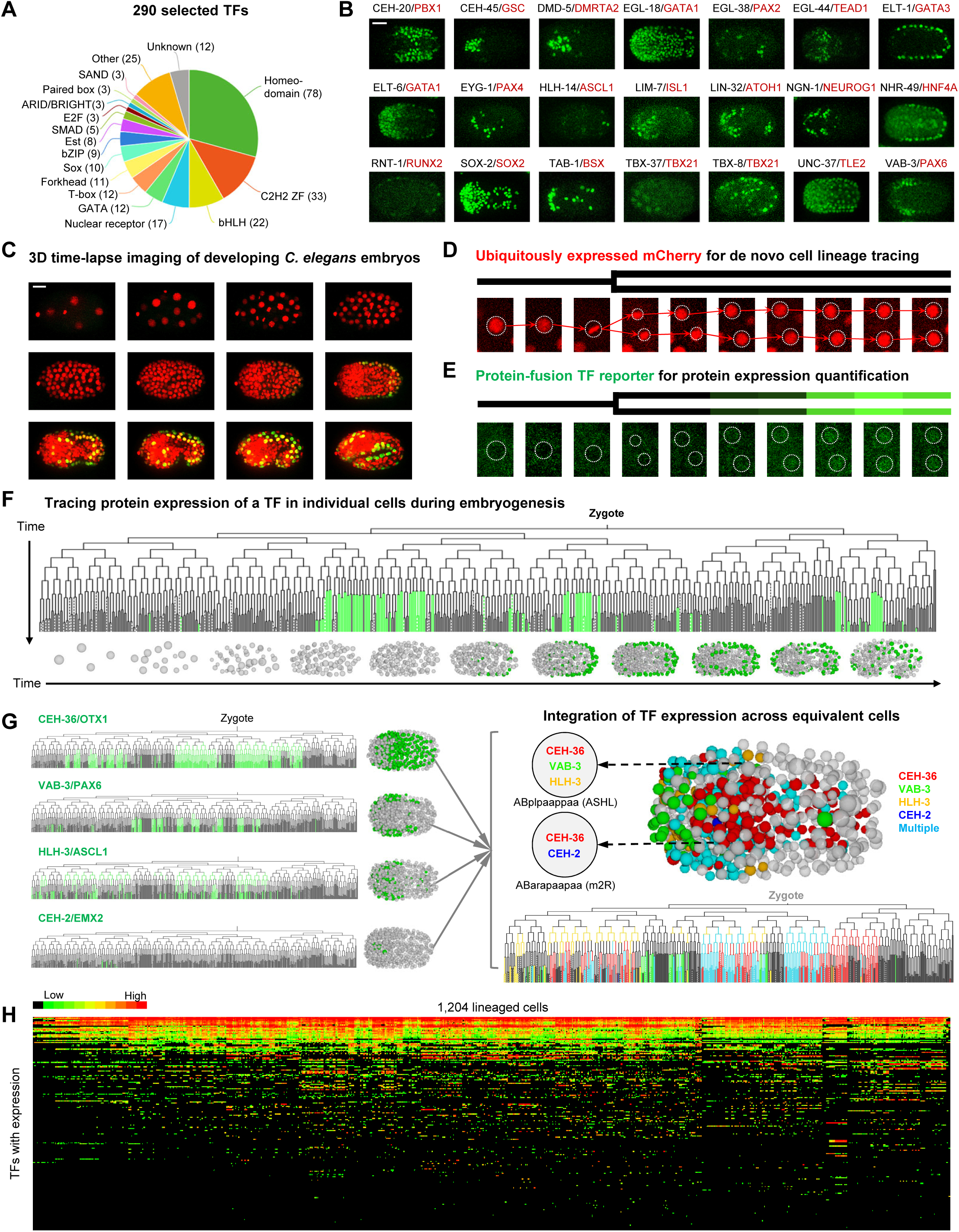
Construction of a quantitative protein expression atlas describing hundreds of TFs in nearly all identity-resolved embryonic cells (A) TF classes based on typical domains. Classes with less than three TFs are not included. (B) Maximum projections of 3D images showing cellular protein expression of representative TFs. Scale bar = 10 µm. (C) 3D time-lapse imaging of embryogenesis. Scale bar = 10 µm. (D) Automated identification (circle) and tracing (arrow) of nuclei. (E) Quantification of TF expression by measuring nucleus green fluorescent intensity over time. (F) Upper: Tree visualization of binary protein expression of a TF in all lineaged cells. Vertical lines indicate cells traced over time and horizontal lines indicate cell divisions. Lower: 3D visualization of TF-expressing cells in the embryo. (G) Schematic of TF atlas construction via integrating the expression of multiple TFs in lineage equivalent cells. Left shows the expression of individual TFs and right shows the integrated result. (H) Heatmap showing quantitative protein expression of TF reporters in 1,204 lineaged cells. See also Figure S1 and Table S1.

To facilitate single-cell protein quantification, we generated 291 dual-fluorescent reporter strains by crossing a ubiquitously-expressed nuclear mCherry reporter into the above strains. For each dual-reporter strain, we performed 3D time-lapse imaging to record over ∼6.5 hours of embryogenesis at 1.25- minute temporal resolution (Figure 1C). Using the nucleus-localized mCherry signal, semi-automated cell identification and tracing were performed to de novo reconstruct entire cell lineages and characterize cell identities up to the bean stage (∼600 cells), when most cells have completed cell differentiation (Figure 1D) (Du et al., 2014; Murray et al., 2008). All traced terminal cells were followed for at least 15 minutes. Simultaneously, GFP/mNeonGreen intensity was measured in each cell at each time point to quantify the dynamic protein expression of the strain’s associated TF (Figure 1E). Thus, the 4D images of reporter expression were digitized into a quantitative measure of protein expression in lineaged cells during embryogenesis (Figure 1F).

We constructed an expression atlas through the cell-by-cell integration of expression for all TFs having reporter strains. *C. elegans* development follows an invariant cell lineage and stereotyped cell divisions to generate a fixed number of progenitor and terminally-differentiated cells (Sulston et al., 1983). Because developmental fate is invariantly associated with lineage history, equivalent cells in different embryos can be straightforwardly identified (Figure 1G). Using the traced cell lineages and cell identities, we integrated the expression of 266 TFs across 1,204 embryonic cells, producing a comprehensive atlas of TF protein expression at high spatiotemporal resolution and coverage (Figure 1H) that covers 90% of all cells generated during embryogenesis.

### A high-quality and informative single-cell protein expression atlas

We performed quality controls to compensate for fluorescence attenuation across depth, thereby ensuring optimized expression quantification, high accuracy of lineage tracing, and reliable cell-to-cell comparison of expression (STAR Methods and Figure S2A-E). We determined expression to be highly reproducible between embryos of the same strain (median r = 0.91, n = 266, Figures 2A and 2B). Expression of the same TF in different reporters was also highly concordant, including between fosmid reporters (median r = 0.91, Figure S2F) and between fosmid and knockin reporters (median r = 0.89, Figure S2G). These results validate the high technical reliability of the reporter strains, lineage tracing results, and expression quantification.

**Figure 2.**
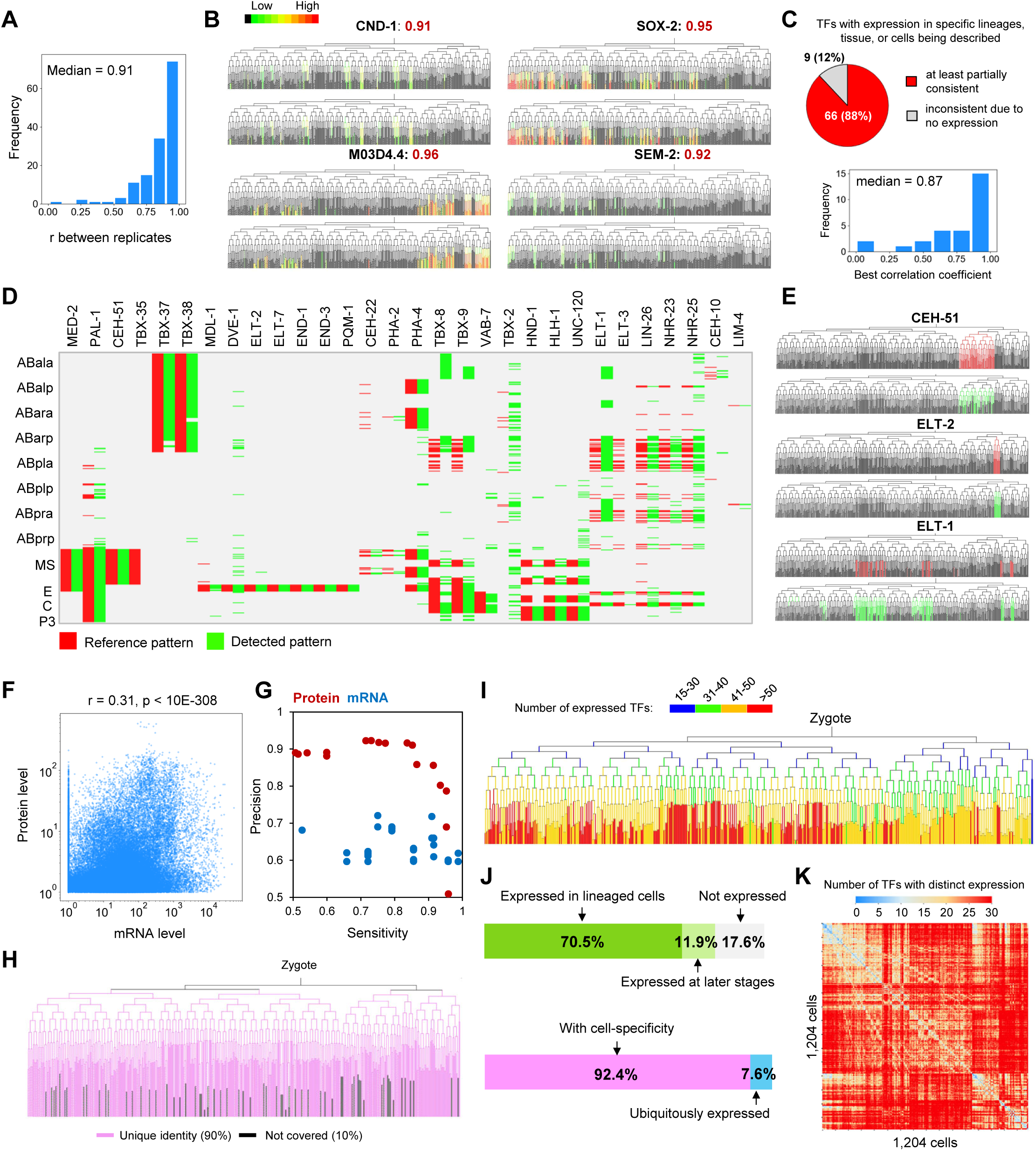
A high-quality and rich dataset of TF expression (A) Distribution of the correlation coefficient for experimental replicates of TF expression assays. (B) Representative examples showing the consistency of cellular TF expression between replicates. (C) Top: Consistency of TF expression patterns between this study and existing literature. Bottom: distribution of correlation coefficient between protein expression of TFs described in this and previous studies. (D) Comparison of the measured TF expression (green) to that reported for 30 TFs selected from the literature (red) in all cell tracks leading to terminal cells. (E) Representative examples showing the consistency of the cellular TF expression described here to that reported in the literature. (F) Correlation between cellular mRNA and protein expression. Statistics: Pearson correlation, n = 213,976. (G) Comparison of the precision and sensitivity of mRNA and protein expression results at different expression cut-offs, using as a benchmark 30 TFs with known cellular expression patterns. (H) Tree visualization of the coverages of analyzed cells and assigned cell identities in this study and one previously published (Packer et al., 2019). (I) Tree visualization of the number of expressed TFs in individual cells. (J) Expression status and specificity of TFs. (K) Heatmap showing the number of TFs exhibiting distinct expression status between all pair-wise cell comparisons. See also Figure S2, Table S2, and Table S3.

We also confirmed that the expression atlas systematically captures known expression patterns. First, we focused on the TFs with sporadic or no expression (expressed in less than five cells) and found that 88% (n = 83) of them were consistent with whole-embryo time-series RNA-seq or scRNA-seq data (Hashimshony et al., 2015; Packer et al., 2019) (Table S3). The inconsistent TFs (n = 9) might have been due to the protein expression being lower than the imaging detection threshold or the protein function being disrupted by fluorescent protein fusion. Next, we compiled a list of TFs from literature for which expression patterns in specific lineages, tissues, or cells were previously described (n = 75). For 88% of the TFs, our expression patterns were at least partially recapitulate the patterns described in the literature (Figure 2C, top, and Table S3). All of the inconsistent cases were on account of our data showing no expression of the TFs. Furthermore, single-cell protein expression of a small number of TFs has been described (Lee et al., 2019; Murray et al., 2012; Sarin et al., 2009; Vidal et al., 2015; Walton et al., 2015); cell-by-cell comparison showed that our data generally correlates well with those described patterns, with a median correlation coefficient of 0.87 (Figure 2C, bottom, and Table S3). In addition, we also more explicitly quantified the accuracy of our data by compiling 30 TFs with known expression restriction and function in specific cells (Table S3). Cell-by-cell comparison revealed that our data achieve a sensitivity and precision of 0.91 and 0.87, respectively (Figures 2D and 2E). These results validate a generally high biological relevance of the atlas.

A valuable lineage-resolved scRNA-seq of *C. elegans* embryogenesis has recently been generated (Packer et al., 2019), raising a question of whether the protein atlas provides additional information. Consistent with previous evaluations (Grun et al., 2014), we found that mRNA levels correlate only weakly with protein levels (r = 0.31, Figure 2F); thus, our dataset provides necessary information on the post-transcriptional regulation and potential function of TFs. Furthermore, using the 30 TFs mentioned above as a benchmark, we found that while mRNA data nicely recapitulate known expression patterns, sensitivity and precision are lower than for the protein data (Figure 2G). Lastly, cell annotation clarity in our data surpasses that of the scRNA-seq dataset. In the scRNA-seq data, cell lineage relationships were determined based on transcriptome similarities between individual sampled cells with the help of a limited number of cell- or lineage-specific marker genes, which approach will inevitably introduce some uncertainty in lineage identity assignments. In contrast, our approach precisely determined cell identities based on direct lineage tracing, assigning unique identity to all lineaged cells (Figure 2H). Together, these results demonstrate that the protein expression atlas provides valuable complementary information on TF expression.

Finally, the protein expression atlas is information-rich. On average, each TF is expressed in 196 cells, and each cell expresses 43 TFs (Figure 2I). Of all TFs evaluated, 82% are expressed during embryogenesis, with 71% in lineaged cells and 12% at later stages (Figure 2J). Significantly, 92% of the expressed TFs exhibit cell-specificity (Figure 2J). Cells tend to express distinct sets of TFs, whereby a given cell can be distinguished from 93% of other cells based on expression of ten or more TFs (Figure 2K). Thus, the atlas provides rich information for defining regulatory states at single-cell resolution.

Collectively, while the atlas described here is TF-centric, the critical role of TFs in cell differentiation, measured protein levels, clarity of cell annotations, high cell coverage, and cell-specificity make it a valuable dataset for elucidating the molecular processes underlying *C. elegans* embryogenesis. In following sections, we demonstrate the usefulness of this dataset in elucidating both TF functions and general principles of the spatiotemporal regulation of cell lineage differentiation.

### A large repertoire of tissue-specific TFs contributes to the ontology-based convergency of regulatory states

The ultimate goal of cell differentiation is to generate sets of specialized terminal cells with distinct functions. Although cell lineages in *C. elegans* have been completely mapped (Sulston et al., 1983), the molecular processes underpinning cell differentiation remain to be fully elucidated. We defined cellular regulatory state (CRS) as the combinatorial expression of TFs in a cell and examined the molecular processes driving differentiation of the totipotent zygote into diverse cell types.

Except for the intestine, all *C. elegans* somatic tissues are multiclonal, containing cells derived from multiple cell lineages (Figure 3A). To determine whether cells making up a tissue type establish a similar CRS, we classified all terminally differentiated cells into five tissue types and used the Uniform Manifold Approximation and Projection (UMAP) algorithm to visualize the distribution of CRSs (McInnes et al., 2018), revealing that cells of each tissue generally clustered together (Figure 3B). Quantitatively, the Jensen-Shannon divergence of CRS between cells of one tissue type (intra-tissue) is significantly lower than that between cells of different tissues (inter-tissue) (Figure S3A). This suggests that, consistent with the recent finding obtained using single-cell transcriptomes (Packer et al., 2019), CRSs generally converge in terminally differentiated cells according to tissue type.

**Figure 3.**
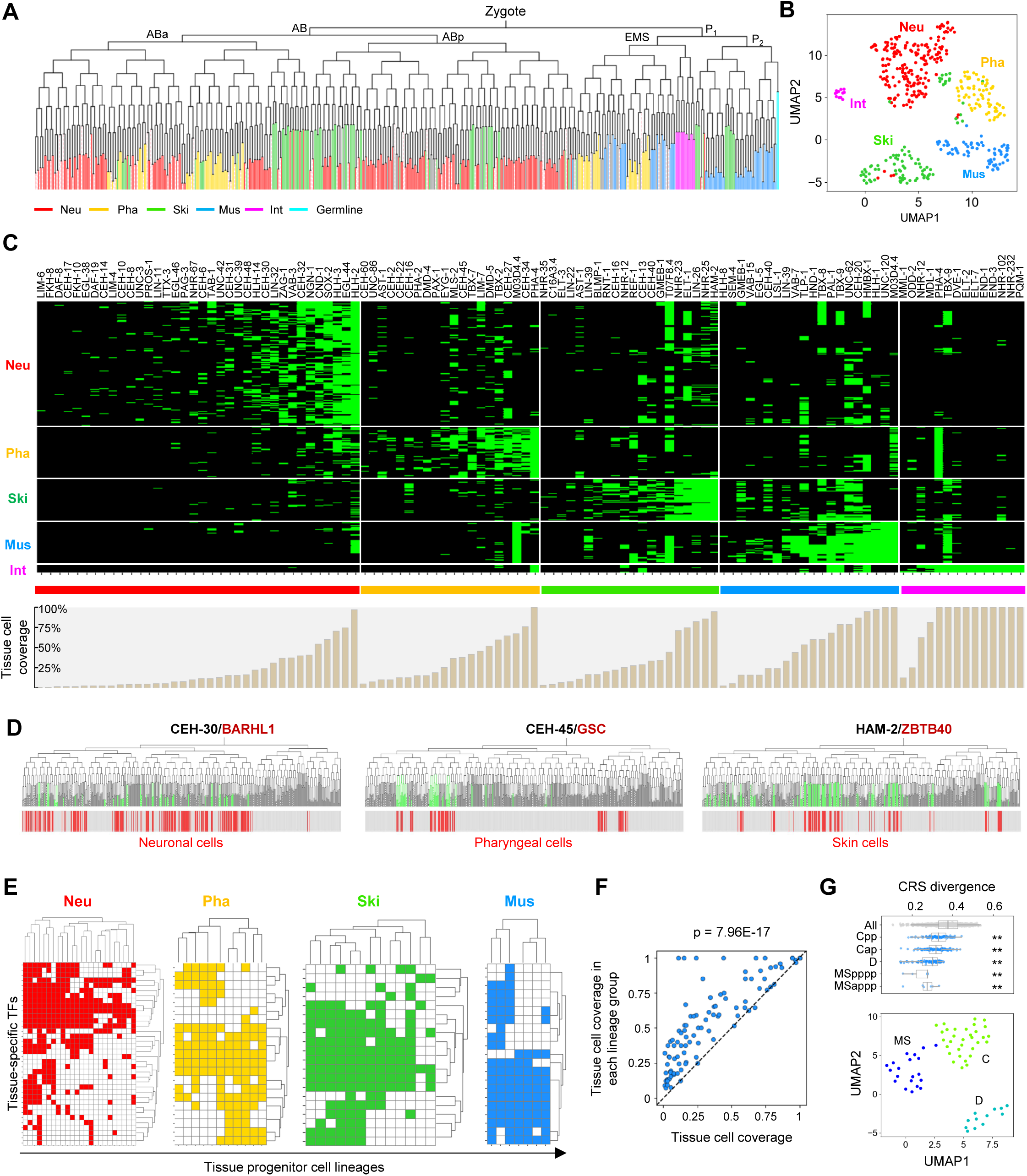
Tissue-specific TFs shape the specificity and heterogeneity of CRS in terminally differentiated cells (A) Distribution of tissue types (colors) across terminal cells. (B) UMAP plot of all terminally differentiated cells (colored by tissue type) based on TF expression. Parameter: min_dist = 0.8, n_neighbors = 10. (C) Top: Heatmap showing the expression of tissue-specific TFs (columns) in all cell tracks leading to terminal cells, ordered by tissue type (rows). Color bars indicate TFs preferentially expressed in corresponding tissues. Bottom: cell coverage of each TF in corresponding tissues. (D) Protein expression of representative tissue-specific TFs in lineaged cells. Bar codes on the bottom highlight cells of specific tissue types. (E) Expression of tissue-specific TFs (rows) in each tissue progenitor cell lineage (columns). (F) Comparison of cell coverage of TFs in corresponding tissues before (x-axis) and after (y-axis) dividing all cells of a tissue type into groups according to their derived progenitor cell lineages. The diagonal line indicates equal cell coverage. Statistics: Wilcoxon signed-rank test. (G) Comparison of CRS divergence between all intra-tissue cells and between intra-tissue cells within each tissue progenitor cell lineage. Statistics: Mann–Whitney U test. ** indicates *p* < 0.01. (H) UMAP plot of MS, C, and D lineage-derived body muscle cells. Parameter: min_dist = 0.8, n_neighbors = 5. See also Figure S3 and Table S4.

To elucidate the molecular basis of this convergency, we identified TFs preferentially expressed in a specific tissue: 101 in total, including 36 for the neuronal system (Neu), 20 for pharynx (Pha), 20 for skin (Ski), 21 for body wall muscle (Mus), and 14 for intestine (Int) (Figure 3C and Table S4). This list overlapped significantly with tissue-specific genes identified by a recent study (Warner et al., 2019), exhibiting 4.4 to 10.3-fold enrichment in different tissues (Figure S3B). Interestingly, many of the tissue-specific TFs identified only in this study are expressed in small subsets of cells (Figure 3C and Figure S3C), demonstrating that single-cell analysis facilitates uncovering TFs with very restricted expression, along with those not previously characterized. For example, we identify CEH-31, BarH like homeobox protein, as preferentially expressed in a subset of neuronal cells; CEH-45, a goosecoid homeobox protein, as pharyngeal-specific; and HAM-2, a zinc finger TF, as skin-specific (Figure 3D). In addition, the cellular expression of tissue-specific TFs is tightly coupled to tissue fate. We selected four TFs, induced fate transformations in associated cell lineages, and observed precise cell-level cut-and-paste of TF expression upon fate transformation (STAR Methods and Figure S3D). Importantly, referencing available genome-wide binding patterns revealed by chromatin immunoprecipitation sequencing (ChIP-seq) experiments for tissue-specific TFs (Kudron et al., 2018), we found their targets to be significantly enriched for genes specifically expressed in the associated tissue (Figure S3E), and for genes whose knockdown elicits phenotypic abnormalities in said tissue (Table S4). These results demonstrate high functional relevance of the identified tissue-specific TFs.

All told, we identified that a significant portion (38%) of TFs exhibit tissue-specific expression that underlies the establishment of tissue-specific CRSs. This not only significantly extends the repertoire of tissue-specific regulatory genes but also facilitates single-cell analysis of the molecular regulation of tissue differentiation.

### Lineage-restricted expression of tissue-specific TFs shapes CRS heterogeneity in terminally differentiated cells

Interestingly, while some tissue-specific TFs are expressed in most cells of a tissue (pan-tissue TFs), a considerable portion are expressed only in a subset (sub-tissue TFs, Figure 3C and Table S4). For example, 80% of neuronal-specific TFs are expressed in less than half of all neuronal cells within the time window assayed here. This is not due to broad classification of type, as identical results were obtained with fine typing (Figure S3F). As most tissues are comprised of cells from multiple lineages, we examined whether lineage origin accounted for restrictions in expression. Specifically, we classified cells into lineage groups which uniformly generate specific tissue types and compared group distributions with sub-tissue TF expression patterns. Indeed, the sub-tissue TFs were preferentially expressed in cells from the same cell lineage group (Figure 3E). The coverage of cells of each tissue by a tissue-specific TF within each lineage group was significantly higher than the overall coverage (Figure 3F).

This lineage-restricted expression of tissue-specific enrichment of TFs explains the widespread intra-tissue CRS heterogeneity (Figure 3B). Although tissue-specific CRSs are generally established in terminal cells, many cells of a given tissue exhibited very high intra-tissue divergences in CRS (Figure S3A). As visualized in the UMAP plot, individual cells of a given tissue type are scattered and form sub-clusters (Figure 3B). This heterogeneity persisted when classifying neuronal and pharyngeal cells into fine groups (Figure S3G), suggesting the heterogeneity is not caused by cell subtypes in a tissue. Interestingly, heterogeneity was significantly reduced when considering one lineage group within a tissue, with cells forming sub-clusters according to origin (Figure 3G and Figure S3H). This suggests the lineage organization of tissues and lineage-restricted expression of tissue-specific TFs contribute to intra-tissue CRS heterogeneity.

Together, both pan- and sub-tissue specific TFs with expression restricted to specific tissue progenitor cell lineages shape the tissue-specificity and heterogeneity of CRSs in terminally differentiated cells. Lineage-restricted expression is a general property of tissue-specific TFs, highlighting a non-negligible influence of origin on a cell’s ultimate state.

### Lineage-specific TFs specify unique states in multipotent progenitor cells before tissue fate specification

Next, we sought to elucidate the molecular basis of the lineage effect on CRS. Interestingly, we found that highly-specific CRSs are efficiently established in early progenitor cells, before tissue fate specification. Pair-wise comparison of TF expression in 162 multipotent progenitor cells revealed that a considerable number of TFs (≥ 5) can be used to distinguish one progenitor cell from most others (≥ 80%, Figure 4A). All told, we identified 84 TFs as preferentially expressed in specific lineages (Figure 4B and Table S4), covering most early multipotent lineages. These included previously-known lineage-specific TFs, such as TBX-37 (ABa), CEH-51 (MS), and PAL-1 (C and D), and uncharacterized ones, such as SPTF-1, ETS-7, and FKH-2 (Figure 4C).

**Figure 4.**
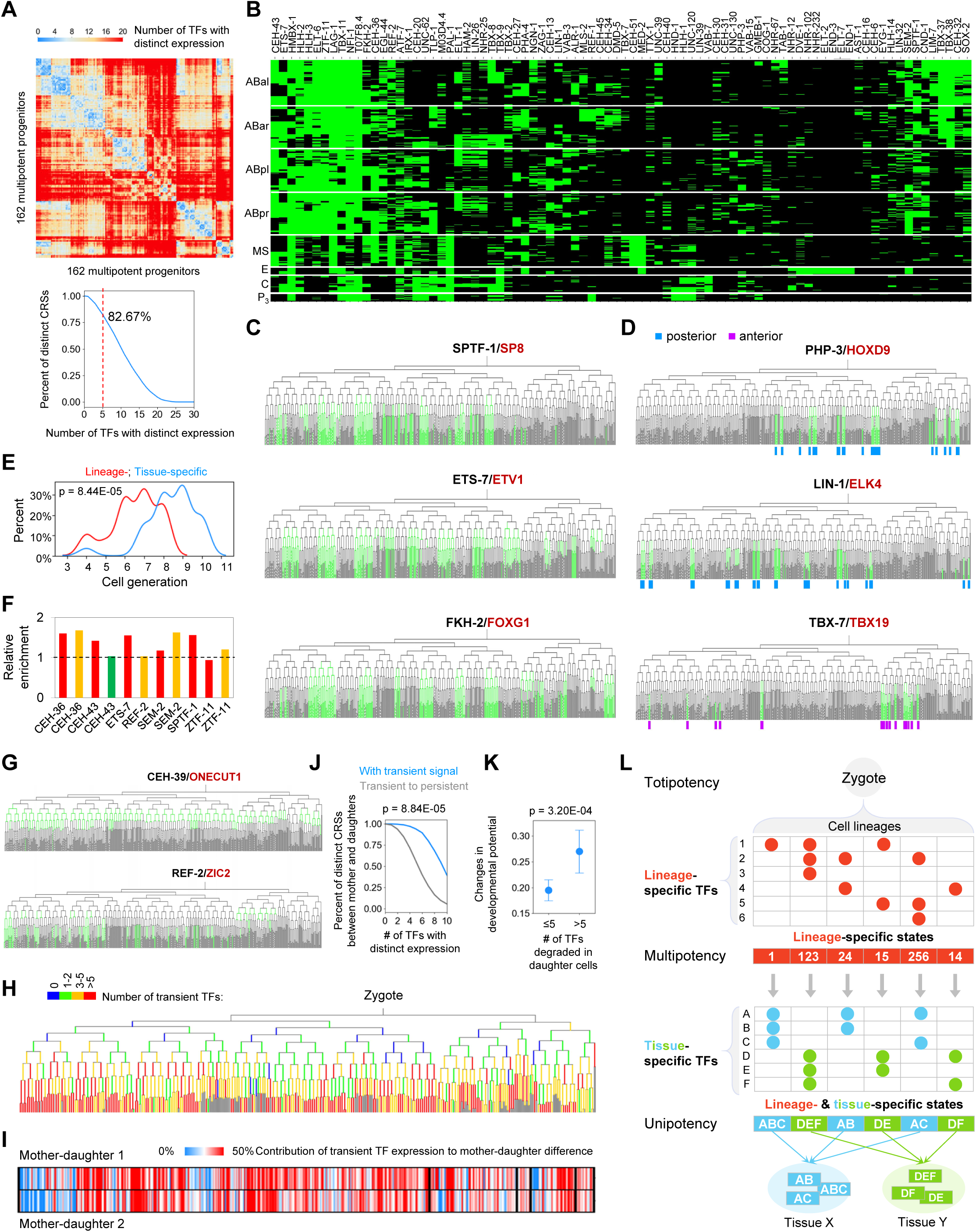
A combinatorial code of lineage-, tissue-, and time-specific TFs for spatiotemporal patterning of CRS (A) Top: Heatmap showing the number of TFs exhibiting differential expression between multipotent progenitors. Bottom: Percent of distinguishable progenitor CSCs under various criteria. (B) Expression of lineage-specific TFs in cell tracks, ordered by lineage. (C-D) Expression of representative lineage-specific (C) and A-P asymmetric (D) TFs in lineaged cells. (E) Expression onset of lineage- and tissue-specific TFs. Statistics: Kolmogorov–Smirnov test. (F) Fold enrichment of tissue-specific genes (Warner et al., 2019) in the targets of lineage-specific TFs. (G) Expression of representative transiently-expressed TFs in lineaged cells. (H) The number of transiently-expressed TFs in each cell. (I) Contribution of transient expression to the net difference in TF expression between mother and daughter cells. (J) Percent of distinguishable mother-daughter CRCs under various criteria before and after considering transient TF expression. Statistics: Wilcoxon signed-rank test. (K) Difference in developmental potential between daughter cells having a small (≤5 TFs) or large (>5 TFs) number of TFs that are degraded in the cells. Statistics: Mann–Whitney U test. (L) Spatiotemporal TF combinatorial code for pattering CRSs during cell lineage differentiation. See the main text. See also Table S4.

A special type of lineage-specificity is anterior-posterior (A-P) asymmetric division, in which distinct CRSs are established repeatedly between two daughter cells (Mizumoto and Sawa, 2007). We identified 16 TFs (including the well-known POP-1/TCF) exhibiting significant A-P biased expression, eight preferentially expressed in posterior lineages (such as PHP-3 and LIN-1) and seven in the anterior lineages (such as TBX-7) (Figure 4D and Table S4). A-P asymmetric division is triggered by the Wnt pathway, and leads to asymmetric localization of regulatory proteins (POP-1/TCF and SYS-1/β-catenin) with the daughter cells having different fates (Mizumoto and Sawa, 2007). Asymmetrically-expressed TFs might transduce the asymmetry of POP-1 and β-catenin into differential CRSs, contributing to the fine patterning of states within lineages.

All told, a large repertoire of lineage-specific TFs specifies distinct CRSs in the early cell lineages. If these TFs also underlie the lineage-restricted expression of tissue-specific TFs, we expected they would tend to target tissue-specific TFs. We found lineage-specific TFs were expressed significantly earlier than tissue-specific TFs (Figure 4E, *p* = 8.44E-05). Furthermore, using available genome-wide binding data, we tested whether lineage-specific TFs tend to target tissue-specific genes. For each lineage-specific TF, we identified the predominant tissue types within the corresponding TF-expressing lineages and determined whether genes specifically expressed in these tissues were enriched for the TF’s targets. With all seven testable exclusive lineage-specific TFs, we found that the frequency of TF target genes specifically expressed in at least one of the predominant tissue types was higher than the expectation (Figure 4F).These results suggest a regulatory hierarchy between lineage- and tissue-specific TFs: Distinct TFs are first expressed in the multipotent progenitor cells of early lineages, activating a cohort of pan-tissue and sub-tissue TFs in corresponding cell lineages.

### Transient expression of TFs increases the temporal diversity of CRS

Interestingly, a large portion (n = 58, 31% of TFs with expression) of TFs exhibit strong transient expression, persisting for only a short time in specific developmental contexts (Table S4). When transient expression detected, the signal typically dampened abruptly, indicating developmentally-programmed degradation. In some cases, such as CEH-39, expression persisted for just two cell generations; in other cases, such as REF-2, multiple rounds of expression were detected (Figure 4G). Frequency calculation in individual cells revealed widespread transient TF expression during embryogenesis, without significant lineage or time bias (Figure 4H).

Time-specific expression of regulatory genes is critical to assigning temporal cellular identity (Kohwi and Doe, 2013). Given that developmental potential is generally more restricted following cell division, we examined the contribution of transient TF expression to distinguishing temporal CRSs between mother and daughter cells in two possible strategies: new expression in the daughter cells, or degradation of mother cell TFs. Intriguingly, the degradation strategy contributed to 31% of the net expression difference, suggesting transient TF expression as a general strategy for distinguishing temporal CRSs (Figure 4I). When transient signals were forced to be persistent in simulation analysis, more than 30% of CRSs that were otherwise distinct between mother and daughter cells (≥ 5 TFs exhibiting differential expression) became indistinguishable (Figure 4J). Finally, we calculated temporal fate transition as the change in differentiation potential from mother to daughter cells, and observed a significantly higher fate transition when the frequency of TF degradation in daughter cells was high (Figure 4K). This suggests that timely removal of TF protein contributes to temporal fate transitions.

In summary, systematic analysis of the cell-specificity of TFs reveals a spatiotemporal combinatorial code specifying CRSs (Figure 4L). First, lineage-specific TFs specify cell-specific CRSs in multipotent progenitor cells, which activate tissue-specific TFs in a highly lineage-dependent manner in corresponding tissue progenitor cells. Expression of both pan-tissue and sub-tissue-specific TFs then drives the establishment of characteristic CRSs in terminal cells according to tissue type while exhibiting widespread lineage-based heterogeneity. During lineage progression, many TFs are also rapidly degraded to distinguish mother-daughter CRS and to increase temporal diversity of the regulatory state. Collectively, these mechanisms achieve a highly spatiotemporal patterning of CRSs.

### A spatiotemporal TF regulatory cascade of cell lineage differentiation

To understand the regulation of differentiation by TFs, we constructed a spatiotemporal regulatory cascade based on inferred functions, cellular expression patterns, co-expression, and regulatory targets. We first deconstructed the cell lineage regime into spatiotemporal regulatory modules and linked TF functions to each module. As regulation of differentiation is highly lineage- and time-dependent, we treated individual progenitor cell lineages as the spatial module of differentiation and further divided each into three temporal modules: upstream of tissue specification, specification or early differentiation, and late/terminal differentiation (Figure 5A and Figure S4A). This divided the differentiation of five tissues into 168 spatiotemporal modules. TFs were assigned to modules based on lineage-, tissue-, and time-specificity of expression (Figure 5B). Lineage-specific TFs were considered to function in the upstream module, with tissue-specific TFs in the specification/early or late/terminal modules according to expression timing.

**Figure 5.**
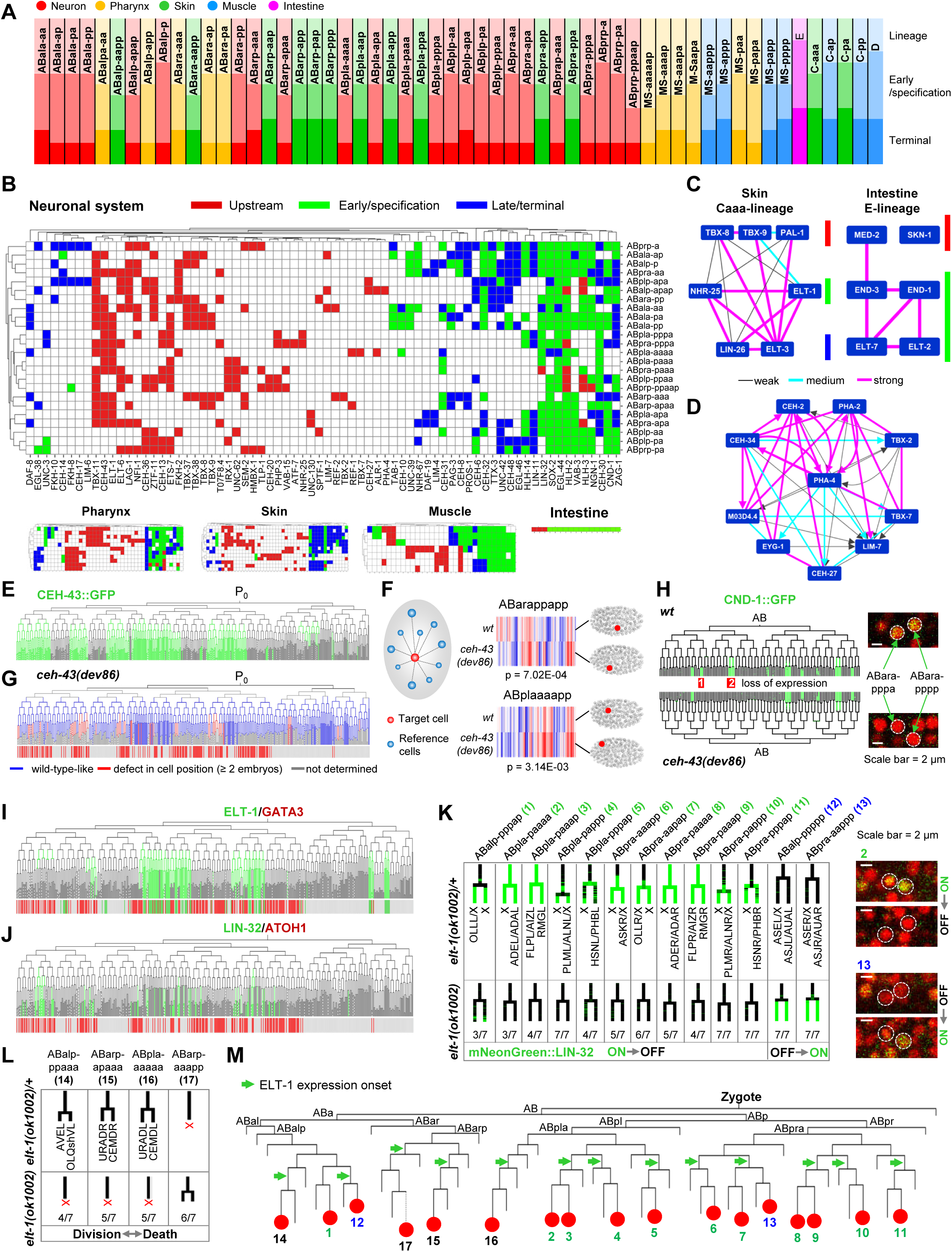
A spatiotemporal TF regulatory cascade of cell lineage differentiation (A) Spatiotemporal modules of cell lineage differentiation. Each column represents the development of a tissue progenitor cell lineage (spatial module), with the name of the founder cell indicated. Each row represents the classification of the regulation of each progenitor cell lineage into three temporal modules (color gradient). (B) The assignment of lineage- and tissue-specific TFs into spatiotemporal regulatory modules of corresponding tissues. Each row is a tissue progenitor cell lineage, and colors indicate the locations of TFs in corresponding temporal modules. (C) Part of the inferred spatiotemporal regulation of skin (Caaa lineage-derived) and intestine (E lineage-derived) differentiation containing the TFs whose function and regulatory relationships have been well established. (D) Expression similarity and regulatory relationship between PHA-4 and all pharyngeal-specific TFs that are expressed in the ABa-derived pharyngeal cells. (E) Protein expression of CEH-43 in lineaged cells. (F) Left: Quantification of relative cell position by measuring the geometric distances between a target cell and all other cells at the same stage (reference cells). Right: Representative examples showing cell position changes in which the heatmap shows the vector of distances for a cell in *wt* and mutant embryos and the 3D rendering graph shows its position in real embryos. Statistics: *t*-test, *p* values were Benjamini-Hochberg corrected. (G) Tree visualization of cell position phenotypes detected in *ceh-43(dev86)* embryos. The barcode on the bottom highlights neuronal cells. (H) Single-cell protein expression of CND-1 in *wt* and *ceh-43(dev86)* embryos. (I-J) Single-cell protein expression of ELT-1 (I) and LIN-32 (J) in lineaged cells. (K) Left: Loss and gain of LIN-32 expression in corresponding cell lineages of *elt-1(ok1002)* embryos. Numbers at the bottom indicate the penetrance of each phenotype. Right: Representative micrographs showing the changes of LIN-32 expression in *elt-1(ok1002)*. (L) Changes in cell lineage patterns in *elt-1(ok1002)* embryos. (M) Summary of ELT-1 function in neurogenesis. See also Figure S4, Table S5, and Table S6.

We then inferred the relationships between TFs in each spatiotemporal regulatory module by linking those with highly similar single-cell expression (Figure S4B); this network represented regulatory relationships inferred from cellular expression. We confirmed the efficiency of this strategy in capturing functional relationships by compiling 30 pairs of TFs known to function in pathways regulating cell differentiation (Table S3). In our dataset, the similarity of expression between these benchmark TF pairs was over four-fold greater than expected (Figures S4C and S4D). As shown in Figure 5C, the inferred spatiotemporal TF regulation of skin and intestine cell differentiation is highly consistent with previously-reported regulatory cascades (Maduro, 2010; Yanai et al., 2008).

We also leveraged available genome-wide binding data (Kudron et al., 2018) as representing experimentally-identified direct regulatory relationships in which one TF directly targets another in the network. For example, we used the ChIP-seq targets of PHA-4, a master regulator of pharynx development (Gaudet and Mango, 2002), to construct a network of PHA-4-regulated TFs (Figure 5D) that not only captured known regulators of pharynx development (e.g. PHA-2, CEH-2, and CEH-22) but also revealed potential new ones (e.g. CEH-27 and EYG-1) (Mango, 2007). We experimentally validated one potential regulator, CEH-27/NKX2, whose highly pharynx-specific expression is similar to that of PHA-4 (Figure S4E). In *pha-4* mutants, CEH-27 protein expression is completely absent in specific ABalp-derived cells that normally differentiate into pharyngeal cells (Figures S4F and S4G). This validates using TF relationships inferred from expression similarity alongside genome binding to identify new genes involved in cell differentiation.

We then constructed a spatiotemporal TF cascade using the same strategy, which contained 5,327 TF relationships in 161 regulatory modules (Table S5), revealing the hierarchy of TF functions in regulating temporal cell differentiation in each tissue progenitor cell lineage. As shown above, the cascade accurately predicted known functions and relationships between TFs in regulating cell differentiation (Figure 5C), and facilitates identifying new TFs with roles in tissue differentiation (Figure 5D), providing a unique opportunity for mechanistic analysis of cell differentiation. Below, we illustrate three case studies to demonstrate that the cellular protein expression atlas and the spatiotemporal cascade of TFs provide valuable guidance for elucidating the molecular regulation of cell lineage differentiation.

### CEH-43 specifically regulates the fates and positions of cells from multiple cell lineages

We identified CEH-43, the *C. elegans* ortholog of Distalless/DLX, as a lineage-specific TF preferentially expressed in multiple lineages (Figure 5E and Figure S4H). CEH-43 is required for the development of anterior hypodermal cells and terminal differentiation of dopaminergic neurons (Aspock and Burglin, 2001; Doitsidou et al., 2013). Given the early onset of CEH-43 expression, we generated a loss of function allele to test whether it regulates early cell fate specification, and found that *ceh-43* mutant allele homozygotes died as embryos (Figure S4I), identical to that reported in a putative null allele (*tm480*) of *ceh-43* (Doitsidou et al., 2013). As significant defects in fate specification and differentiation usually affect the 3D position of embryonic cells (Schnabel et al., 2006), we systematically quantified cell position phenotypes in *ceh-43* mutant embryos using an established method (Li et al., 2019) (Figure 5F). Indeed, the relative 3D positions of many early embryonic cells differed significantly from *wild type* (*wt*) (Figure 5F and Table S6); furthermore, the vast majority of affected cells were CEH-43-expressing cells (Figure 5G). The set of cells with abnormal positions included previously-described skin (ABplaaa lineage) cells with developmental defects. We found many ABplaaa lineage-derived cells that normally differentiate into skin cells exhibited extra divisions (Figure S4J), confirming a requirement of *ceh-43* for normal lineage differentiation of skin cells. Interestingly, we also identified progenitors of neuronal cells as having position defects (Figure 5G), suggesting CEH-43 also extensively regulates early neuronal specification. Expression of CND-1/NEUROD1 (Hallam et al., 2000), a reporter of early neurogenesis, was completely abolished in the ABalppap and ABarapp lineages of *ceh-43* mutants (Figure 5H and Figure S4K). Together, these results reveal an indispensable role of CEH-43 in multiple lineage differentiation events and uncover its essential role during early neurogenesis.

### ELT-1/GATA3, a well-known skin fate specifier, functions in neuronal fate specification

ELT-1, a GATA family TF, is a master regulator of skin differentiation, both required and sufficient to drive the specification and differentiation of skin fate (Gilleard and McGhee, 2001; Page et al., 1997). Interestingly, as predicted by the TF cascade, ELT-1 also regulates the differentiation of ABalap-, ABalpp-, ABarp, ABpla-, and ABpra-derived neurons (Figure S4L). ELT-1 is specifically expressed in cells from those lineages (Figure 5I), and ChIP-Seq further showed that ELT-1 binds at the promoters of several well-known neuronal-specific TFs, such as LIN-32/ATOH1, CND-1/NEUROD1, and EGL-44/TEAD1 (Figure S4M). These results suggest an uncharacterized function of ELT-1 in neuronal differentiation.

Among ELT-1 target genes, LIN-32 is of particular interest, being a proneural TF required for specifying the fate of many neuronal cells (Zhao and Emmons, 1995) and widely expressed in many neuronal cells from the above five lineages (Figure 5J). We found its cellular expression to be significantly affected in *elt-1* mutant embryos, with loss in LIN-32-positive neuronal cells and gain in normally LIN-32-negative neurons (Figure 5K, Figure S4N, and Table S6). This confirms that ELT-1 regulates LIN-32 expression in many developmental contexts. Furthermore, *elt-1* mutants exhibited abnormal lineage patterns for several neuronal progenitor cells; for example, programmed cell death was induced in several neuronal progenitor cells that would otherwise divide, and vice versa (Figure 5L).

These results reveal an indispensable role of ELT-1, a well-characterized skin fate specifier, in neurogenesis (Figure 5M). Interestingly, ELT-1 is transiently expressed in ABalap and ABalppp lineages (Figure 5I), wherein alteration of LIN-32 expression was detected; thus, early transient expression could be essential in regulating later cell differentiation. Finally, as ELT-1 is widely expressed in multiple cell lineages that produce both skin and neuronal cells and is required for differentiation of both tissues, we propose that ELT-1 functions as a general lineage specifier rather than a skin-specific regulator.

### M03D4.4 is a pan-muscle TF that guides converging muscle differentiation in the body wall and pharynx

We also identified a TF that is specifically expressed in and participates in the converge of differentiation of the same cell type across different body locations. As predicted by the spatiotemporal module (Figure S5A), M03D4.4, an uncharacterized C2H2-type zinc finger TF, functions in the early and late differentiation of many pharyngeal and body muscle cell lineages. The cellular expression of M03D4.4 is highly specific to muscle cells and their precursors, including the pharyngeal muscle, body muscle, and other muscle cells (Figure 6A), with 88.6% of M03D4.4-expressing terminal cells being muscle cells or sister cells of muscle, and the TF detected in 94% of all muscle cells (Figure 6B). Furthermore, fate switching experiments showed a gain of M03D4.4 expression when transforming other cell types into pharyngeal or body muscle (Figure S5B), confirming M03D4.4 expression is tightly coupled to muscle fates.

**Figure 6.**
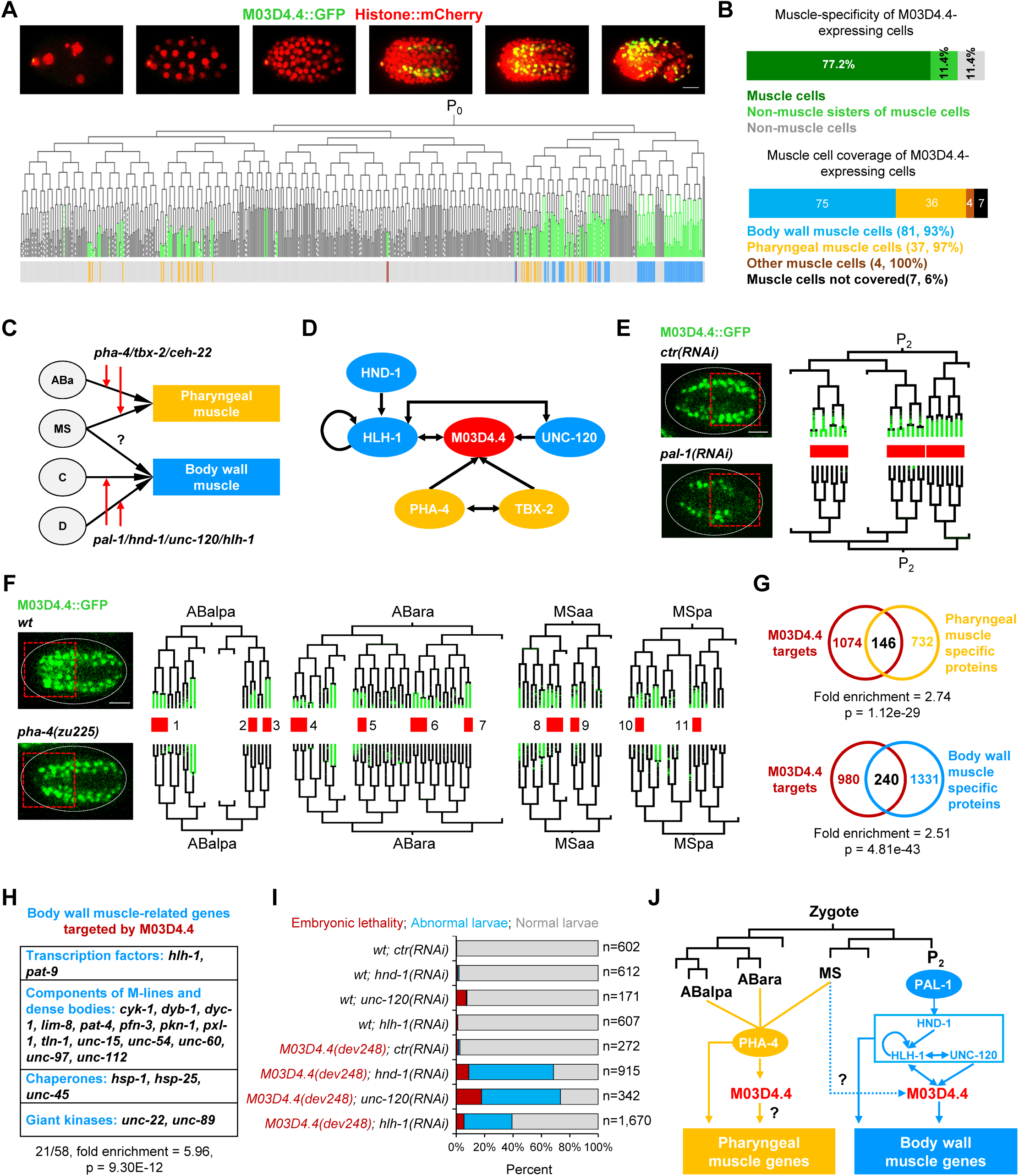
*M03D4.4* encodes a pan-muscle-specific TF responsible for converging muscle cell differentiation in different tissues (A) Top: Maximum projection of 3D images showing cellular protein expression of M03D4.4 at characteristic embryonic stages. Scale bar = 10 µm. Bottom: Expression of M03D4.4 in lineaged cells. The barcode on the bottom highlights pharyngeal (orange), body (blue), and other types of muscle cells (brown). (B) Pan-muscle specificity of M03D4.4. (C) Known pathways underlying fate specification of pharyngeal and body muscle cells derived from different cell lineages. (D) Inferred regulatory relationship between *M03D4.4* and other regulatory genes. (E) Expression of M03D4.4 in cells from the body muscle cell lineages of *wt* and *pal-1(RNAi)* embryos. (F) Expression of M03D4.4 in cells from the pharyngeal muscle lineages of *wt* and *pha-4(zu225)* embryos. (G) Enrichment of proteins preferentially expressed in pharyngeal and body muscle cells (Reinke et al., 2017) in the ChIP-seq targets of M03D4.4. Statistics: Hypergeometric test. (H) Enrichment of genes involved in body muscle development and function (Gieseler et al., 2017) in the ChIP-seq targets of M03D4.4. Statistics: Hypergeometric test. (I) Genetic interactions between *M03D4.4* and regulatory genes of body muscle differentiation. (J) The proposed function of M03D4.4 in the regulatory pathways governing muscle cell differentiation. See also Figure S5 and Table S6.

Next, we demonstrated that the regulatory pathways of pharyngeal and body muscle differentiation converge on M03D4.4. Specification of pharyngeal and body muscle cells is regulated by distinct pathways (Figure 6C); *pha-4/FOXA* functions as an organ identity gene to specify all pharyngeal muscle fates (Gaudet and Mango, 2002), while *pal-1/CDX1/caudal* specifies body muscle fate by activating three redundant TFs, *hnd-1/PTF1A*, *hlh-1/MYOD,* and *unc-120/SRF* (Fukushige et al., 2006; Mango, 2007). Expression of M03D4.4 is highly similar to that of TFs functioning in body (Figure S5C) and pharyngeal (Figure S5D) muscle differentiation. In addition, ChIP-seq data showed that *M03D4.4* is co-targeted by TFs essential for pharynx and body muscle differentiation (Figure 6D), being a direct target of two body muscle regulatory TFs, HLH-1 and UNC-120, and two pharyngeal muscle regulatory TFs, PHA-4 and TBX-2. Importantly, expression of M03D4.4 is dependent on genes in both pathways; knockdown of *pal- 1* caused complete loss of M03D4.4 expression in all C and D lineage-derived cells that normally differentiate into body wall muscle (Figure 6E, Figure S5E, and Table S6), while *pha-4* mutation abolished M03D4.4 expression in many cells that normally differentiate into pharyngeal muscle (Figure 6F, Figure S5F, and Table S6).

Finally, we provide evidence that M03D4.4 regulates differentiation of both body and pharyngeal muscle cells. ChIP-seq data revealed M03D4.4 targets as significantly enriched for proteins specifically expressed in body and pharyngeal muscle cells (Reinke et al., 2017) (Figure 6G), genes important to body muscle development and structure (Gieseler et al., 2017) (Figure 6H), and genes whose targeting by RNAi elicits muscle abnormalities (Figure S5G). We created large-deletion alleles of *M03D4.4* (Figure S5H) and found that while M03D4.4 loss of function alone does not lead to apparent developmental defects, the gene functions synergistically with *hnd-1*, *hlh-1*, and *unc-120* to regulate body muscle development, affecting a large portion of the animal (39% to 73%) and culminating in synthetic embryonic lethality and larval arrest (Figure 6I). Meanwhile, analysis of genetic interaction between *M03D4.4* and pharynx regulators (*pha-4*, *pha-2*, *ceh-22*) failed to identify synthetic phenotypes (Figure S5J and data not shown), leaving its function in pharyngeal muscle development to be elucidated.

Together, analysis of cellular expression, direct targets, and regulatory and genetic relationships revealed that M03D4.4 functions as a pan-muscle TF to direct convergence of muscle differentiation programs initiated by distinct upstream regulators in the body wall and pharynx (Figure 6J).

### Dynamics of CRS reveal a winding trajectory of cell states during differentiation

A fundamental question in developmental and systems biology concerns how CRSs change along the developmental trajectories leading to terminally differentiated cells. The precisely-mapped developmental trajectories of each *C. elegans* cell and the high-resolution spatiotemporal atlas of TF expression described here provide a unique opportunity for addressing this question. We thus quantified the dynamics of CRSs during cell differentiation and revealed general properties of the regulatory landscape.

As a cell progresses through a lineage program, CRSs could theoretically diversify in two dimensions: across cells from different lineages at a given stage (cross-lineage distance) and across cells of different generations within the same cell track (cross-generation distance) (Figure 7A). If CRS changes continuously, we expected the divergence would increase with these two types of developmental distance. If switch-like, changes in CRS would be discrete following an increase of developmental distance. We quantified CRS divergence (difference in TF expression) between cells as a function of developmental distance (Figure 7A) and found that it increased with both cross-generation distance (Figure 7B) and cross-lineage distance (Figure 7C and Figure S6A). This indicates that CRS continues to change during cell lineage differentiation, lacking an apparent quiescent stage.

**Figure 7.**
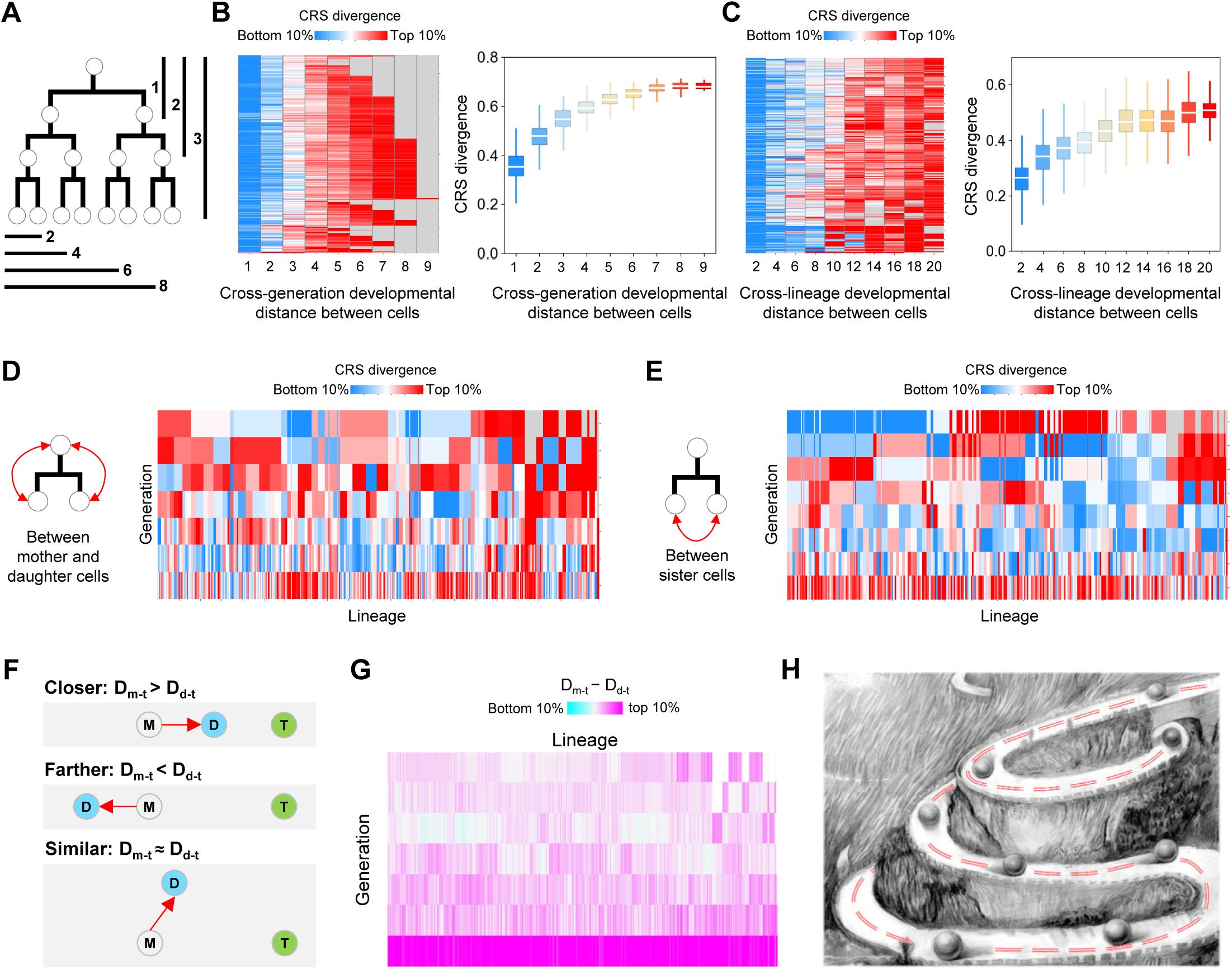
General properties of CRS dynamics during cell lineage progression (A) Definition of the developmental distance between cells. Cross-generation distance is measured as the number of cell divisions from the earlier to the later cell on a cell track. Cross-lineage distance is quantified as the total number of cell divisions from the lowest common ancestor of the two cells to each of the cells at a given developmental stage. (B) Left: Each row shows the change in CRS divergence as a function of cross-generation distance (columns) between all cells on a cell track. Right: Change in CRS divergence as a function of cross-generation distance across all cell tracks. (C) Left: Each row shows the CRS divergence of a cell at the 600-cell stage relative to all other cells at given cross-lineage distances (columns). Right: Change in CRS divergence as a function of cross-generation distance across all cells. (D-E) Heatmap showing CRS divergence between all mother and daughter cells (D) and between two sister cells (E) following all cell divisions. Cell divisions are organized first by generation and then by lineage. (F) Schematic of three scenarios in which the transition of CRS from mother (M) to daughter (D) cell causes CRS divergence of the daughter from the terminal (T) cell to be closer, farther, or similar to that between the mother and the terminal cell. (G) Heatmap showing the difference between D_m-t_ and D_d-t_ following all cell divisions. (H) Detour model of CRS change during cell differentiation. See also Figure S6.

Next, we examined whether CRS dynamics at various developmental stages proceed with constant or varying speed. Comparison of divergence between mother and daughter cells at different developmental stages showed that CRS changes are highly variable in magnitude across time and lineage (Figure 7D), with a similar trend in the divergence between daughter cells across time and lineage (Figure 7E). This suggests that although CRS changes continuously during embryogenesis, the speed of change is highly time- and lineage-dependent.

We further analyzed the directionality of CRS transitions along each developmental trajectory to determine whether CRS progressively approaches the final state of the terminally differentiated cell. In theory, there exist three scenarios: each change in CRS could bring it closer to, farther from, or equally similar to the terminal CRS (Figure 7F). We measured CRS divergence for each mother-daughter progenitor cell pair on a given cell track leading to a terminal cell, determining divergences between mother and terminal cells (D_m-t_) and between daughter progenitor and terminal cells (D_d-t_). Plotting differences between D_m-t_ and D_d-t_ revealed that except for the last round of cell division, D_m-t_ is comparable to D_d-t_ (Figure 7G). This suggests that transitions of CRS between mother and daughter cells generally do not move towards the terminal CRS. We further categorized divergence scores as whether the daughter progenitor cell was closer to the corresponding terminal cell than was the mother cell (Figure S6B), revealing that the last round of cell divisions is highly enriched for CRS transitions that drive the cell toward the terminal state (3.28 fold-enrichment, *p* = 9.52e-280). Following earlier cell divisions, however, CRS transitions do not considerably affect divergence from the terminal cell. Interestingly, we did not observe any cases in which the transition of CRS from mother to daughter cell caused the cell away from the terminal state, although such is theoretically possible. Further investigation of the correlation of CRS divergence between mother and daughter (D_m-d_) cells with the difference between D_m- t_ and D_d-t_ revealed no correlation for most cases (Figures S6C and S6D). These results suggest that a cell’s trajectory from initial to terminal states undergoes extensive detours in which mother-to-daughter CRS transitions are generally not directional towards the terminal state, except upon the last round of cell division (Figure 7H).

Together, systems-level analysis reveals that CRS changes continuously but at a highly variable speed across cells from different lineages and generations. Importantly, changes in CRS do not necessarily drive toward the terminally differentiated state but take a winding path, indicating that the landscape guiding cell differentiation is not straightforward.

## DISCUSSION

### A single-cell protein expression atlas of regulatory genes during lineage differentiation

Mapping the protein expression atlas of regulatory genes is an important step in elucidating the molecular basis of biological processes (Thul et al., 2017). While generating massive single-cell transcriptome data has become routine and straightforward, high-throughput quantification of protein expression at single-cell level remains challenging (Marx, 2019). Recently, single-cell proteomics approaches have been applied to map dynamic cellular protein expression during human erythropoiesis (Palii et al., 2019) and chick hair-cell development (Zhu et al., 2019). However, systematically mapping single-cell protein expression in all lineage-resolved cells during embryogenesis has not been feasible. Leveraging our imaging-based single-cell analysis method, we have mapped a comprehensive *C. elegans* protein expression atlas containing the dynamic expression of hundreds of conserved TFs in nearly all identity-resolved cells during embryogenesis (Figure 1). As protein levels generally cannot be accurately predicted by mRNA expression (Figure 2F) and are in many cases the more reliable representation of gene function (Buszczak et al., 2014), this atlas provides a valuable resource on metazoan embryogenesis that complements the recently generated single-cell mRNA expression atlas (Packer et al., 2019). In this study, we traced cell lineage until a stage when most of the terminal embryonic cells had existed for at least 15 minutes (bean stage); however, due to significant technical challenges in tracing cells in late embryos, these cells were not further traced to a terminally differentiated stage. We noticed from the images that a small number of TFs not expressed in the traced terminal cell within the analyzed time window were indeed expressed later and <u>vice versa</u>, resulting in potential false-negatives or –positives for TF expression in some terminal cells (n = 16, Figure S7). Nevertheless, the combination of high cell coverage, resolved lineage identity, high temporal resolution, and highly cell-specific patterns in the protein atlas (Figures 1-4) provides a unique opportunity to investigate protein regulation and function during embryogenesis.

### A spatiotemporal combinatorial code of TFs drives cell differentiation along lineages

We identified over 70% of expressed TFs as exhibiting lineage-, tissue-, or time-specificity, revealing a TF combinatorial code key to patterning CRS during cell differentiation (Figures 3-4). Interestingly, CRSs are highly patterned in early progenitor cells having multipotency (able to produce two or more tissue types) by means of a cohort of lineage-specific TFs (Figure 4A). Hence, the CRSs of progenitor cells from different lineages are intrinsically different, allowing initiation of lineage-specific programs to drive tissue differentiation. Consistent with this, a large fraction of tissue-specific TFs are expressed in subsets of tissue cells in a lineage-restricted manner (Figures 3C and 3E). Intriguingly, while cells of one tissue type that originate from different progenitor cells have CRSs that converge during late embryogenesis, lineage-specific patterning persists in all tissues (Figure 3G and Figure S3H). Thus, lineage organization contributes to intra-tissue heterogeneity. This phenomenon is not specific to *C. elegans*, whose development follows an invariant lineage; studies of human hematopoietic differentiation have identified a lineage bias in cell subtypes, which follow distinct developmental trajectories (Buenrostro et al., 2018; Velten et al., 2017).

Another intriguing finding is that within a tissue type, the subsets of cells expressing tissue-specific TFs overlap highly (Figure 3E); this suggests that many TFs synergistically regulate tissue differentiation in a lineage-dependent manner, and partially explains the paradox that although TFs are critical regulators of differentiation, loss of function often does not produce defects. Not hinging on one or a few TFs increases the robustness of tissue differentiation, a pattern best illustrated in the regulation of neurogenesis. It is well established that terminally differentiated neurons of different types display highly different states regulated by a collection of terminal selector transcription factors that are expressed and function in a combinatorial manner (Hobert, 2016). Here, we show that the expression of neuronal-specific TFs in progenitor cells is also highly redundant and overlapping.

In addition, we revealed that many TFs exhibit strong transient expression that increases the temporal diversity of CRSs (Figures 4G-4K). Timely degradation of specific TFs is critical in driving the transition of differentiated states; for example, turnover of SKN-1, a master TF for *C. elegans* endomesoderm, is required for restricting developmental potential from endomesoderm to mesoderm (Du et al., 2015a; Du et al., 2014). Here, we show that transient expression of ELT-1 in neuronal cell lineages is essential for specifying early neuronal fate (Figures 5I and 5K). Extensive studies in *Drosophila* have established that temporal-specific expression and degradation of several TFs contributes to temporal patterning and expands neural diversity (Doe, 2017; Miyares and Lee, 2019). Our finding of a large number of transiently expressed TFs suggests that developmentally programmed degradation is a general paradigm regulating differentiation. More broadly, this single-cell analysis of combinatorial TF expression reveals general principles on efficiently patterning CRS in a metazoan embryo.

### A winding landscape of cellular state transition during cell differentiation

Systems-level analysis of CRS dynamics over the course of lineage progression revealed important kinetics of *in vivo* cell differentiation; namely, CRS progression is not directional until late embryogenesis (Figure 7). This suggests that changes in regulatory signal (TF expression) do not immediately drive the cell toward its ultimate state. Significantly, the opposite scenario in which changes in TF expression and CRS cause cells to diverge greatly from their final destinations is systematically repressed (Figure 7G), suggesting that while not straightforward, the trajectory of cell differentiation is highly optimized. One interesting possibility is that seemingly ‘unnecessary’ changes in CRS confer upon the cells a state poised for rapid and directional transition upon terminal differentiation.

### Expression-guided mechanistic analysis of TF function and molecular regulation of cell differentiation

Finally, we have provided three examples reflecting important guidance from the spatiotemporal expression of TFs in revealing TF functions and the regulation of differentiation (Figures 5-6). First, having determined single-cell expression patterns and specificity, potential action sites and functions become readily predictable and testable. Second, similarity in cellular expression between new and known TFs enables direct testing of regulatory relationships and thus pathway placement of the new TF. The large number of protein-fusion reporters collected here facilitates systematic analysis of TF regulatory relationships through perturbation experiments. Third, determination of single-cell expression patterns allows rational prediction and testing of genetic interactions and synergy between TFs. Many tissue-specific TFs are expressed redundantly in single cells and may function synergistically in developmental contexts; we can now more reasonably infer and test potential synthetic effects. If genetic redundancy is widespread, this expression-guided strategy would be more efficient than genetic screening in discovering regulators of complex tissue differentiation. Lastly, we have collected and generated protein-fusion reporters (GFP or mNeonGreen) for over 260 TFs (86 new endogenous protein fusion reporters in this work), strains expressing which can be directly used in ChIP-seq experiments to identify potential targets. With the single-cell ChIP-seq techniques developed recently (Ai et al., 2019; Wang et al., 2019) and the single-cell expression patterns resolved here, building a cell-resolution molecular circuit of *in vivo* lineage differentiation is imaginable.

## ACKNOWLEDGMENTS

We thank Dr. Meng-Qiu Dong, National Institute of Biological Sciences for providing CRISPR/Cas9 reagents; and Caenorhabditis Genetics Center (CGC) for providing some strains. This work was supported by grants from the “Strategic Priority Research Program” of the Chinese Academy of Sciences to Z.D. (XDB19000000), the National Natural Science Foundation of China to Z.D. (31722035 and 31771598) and to X.M. (31900578), and the State Key Laboratory of Molecular Developmental Biology, China.

## AUTHOR CONTRIBUTIONS

Z.D., X.M. and Z.Z. conceived the project and designed the study; X.M., L.X., Z.D., W.X., Y.W., Y.Z., and G.W. conducted the experiments and generated the data; Z.Z. and Z.D. performed the data analysis; Z.D. wrote the manuscript with input from all authors.

## STAR METHODS

### RESOURCE AVAILABILITY

#### Lead Contact

Further information and requests for resources and reagents should be directed to and will be fulfilled by the Lead Contact, Zhuo Du

#### Materials Availability

Plasmids and transgenic *C. elegans* strains generated in this study will be made available on request to the lead contact.

#### Data and Code Availability

All data support this study are provided in Supplementary Tables.

### EXPERIMENTAL MODEL AND SUBJECT DETAILS

#### *C. elegans* strains and culture

All *C. elegans* strains used in this study and their genotypes are provided in Table S1. Some strains were obtained from the *Caenorhabditis* Genetics Center. Unless otherwise noted, all strains were cultured at 21 °C in incubators.

### METHOD DETAILS

#### Selection of TFs

We selected 290 TFs as our targets from a list of 639 potential TFs encoded in the *C. elegans* genome, which list included 528 previously defined high-confidence TFs (Reece-Hoyes et al., 2005) and 111 additional TFs compiled from the literature. Based on OrthoList 2, a compendium of *C. elegans* genes with human orthologs (Kim et al., 2018), WormBase gene annotations (Harris et al., 2020), and the Expression Patterns in *C. elegans* (EPIC) database (Murray et al., 2012), we selected 273 TFs that have human homologs and are likely not ubiquitously expressed. Furthermore, an additional 17 TFs that are not conserved but of potential interest in *C. elegans* cell lineage differentiation, such as *med-1/2*, *lin-26*, and *ref-1*, were also added to the final list.

#### Generation of dual-fluorescent reporters of TFs

We drew upon three sources to generate 291 dual-fluorescent reporters for 266 TFs (Table S1). First, fosmid-based protein-fusion GFP reporters have been generated for 185 of the selected TFs by the ModENCODE project (Araya et al., 2014; Sarov et al., 2012). We crossed these transgenic reporters into a lineage strain (RW10226 or JIM113) in which nucleus-localized mCherry is ubiquitously expressed in all nuclei, thereby enabling cell identification and lineage tracing; we thus generated 202 dual-fluorescent reporters. GFP insertion sites for all fosmid-based reporters were examined by PCR and sequencing using a universal primer targeting the GFP sequence and a gene-specific primer. If PCR failed, whole-genome sequencing was performed to identify the GFP insertion site to determine which gene the GFP tagged. If the GFP tagged another TF also present in our list (n = 8), the strain was kept with the gene annotation corrected. If the GFP was fused to a non-TF gene (n = 5), the reporter was discarded (Table S1). It should be noted that this discrepancy does not necessarily mean the original strains in the Caenorhabditis Genetics Center (CGC) stock are incorrect; it could have arisen during strain labeling, handling, or shipping. In total, 197 reporters for 180 TFs belonged to this category. Second, protein-fusion knockin reporters have been previously generated for eight TFs, including *ast-1*, *bnc-1*, *end-1*, *elt-2*, *lin-15B*, *unc-3*, *unc-30*, and *unc-55* (Kerk et al., 2017; Li et al., 2019; Lloret-Fernandez et al., 2018; Paix et al., 2014; Yu et al., 2017). These transgenes were similarly verified by PCR and then crossed into the lineage strains. Third, for the rest of TFs without a protein fusion reporter, we took the lineage strain JIM113 as genetic background (Walton et al., 2015) and performed CRISPR/Cas9-mediated gene knockin to fuse the coding sequence of the fluorescent protein into the N- or C- terminus of the endogenous target TF. We successfully generated 94 dual-fluorescent reporters belonging to this category. Detailed information on how the dual-fluorescent reporters of TFs were sourced and generated is provided in Figure S1D.

#### CRISPR/Cas9-mediated mNeonGreen/GFP knockin

To visualize endogenous protein expression of a TF, we performed CRISPR/Cas9-mediated knockin to insert a codon-optimized coding sequence of the mNeonGreen into the N- or C-terminal of the genomic locus of the TF (Hostettler et al., 2017). First, two single-guide RNAs (sgRNAs) targeting the regions adjacent to the start (N-terminal fusion) or stop (C-terminal fusion) codons of target TFs were designed using the CRISPR Design Tool (http://zlab.bio/guide-design-resources) (Table S1). The synthesized sgRNAs were cloned into the pDD162 plasmid (Addgene plasmid #47549), which contains an expression cassette for Cas9, using circular PCR and following a previously established protocol (Paix et al., 2014). Second, a homologous recombination (HR) repair template plasmid was constructed for each TF, consisting of the mNeonGreen sequence (coding region plus 6x His tag and 6x flexible glycine) flanked by 5’ and 3’ arms (∼800-1000 bp each) homologous to the endogenous gene; these were synthesized and cloned into the pPD95.77 vector (Addgene plasmid #37465). To prevent the cutting of HR templates by Cas9, point mutations were introduced into the HR template sequences that were targeted by the sgRNAs; synonymous if the target site was located in a coding region, or a C-to-G or A-to-T mutation if in a non-coding region. Complete sequences of HR templates are provided in Table S1. Third, for each TF knockin, the above plasmids plus an editing reporter plasmid pRF4 (inducing roller phenotype) were purified with the PureLink PCR Micro kit (Invitrogen, K310050), and injected into the worm gonads at a final concentration of 50 ng/µL each. F1 animals with the roller phenotype were singled onto a new plate and allowed to generate sufficient offspring before PCR detection of the knockin event using primers targeting the mNeonGreen sequence. Animals with homozygous knockin were identified in the F2 population by PCR.

#### CRISPR/Cas9-mediated gene mutation

Except for the HR template plasmid, which is not needed for generating gene mutations, the procedure for CRISPR/Cas9-mediated gene mutation was identical to the knockin experiments. Multiple sgRNAs were designed for a gene of interest to ensure high efficacy, and multiple mutant alleles were obtained. All mutants were outcrossed at least once before being used for phenotypic analysis. Sequences for all sgRNAs are provided in Table S1.

#### Embryo mounting and 3D time-lapse imaging

Embryo preparation and mounting were performed according to a previously described procedure with minor modifications (Bao and Murray, 2011). First, young adult worms were cut open to release early embryos in a droplet of M9 buffer on a Multitest slide (MP Biomedicals). Second, embryos at 2- to 4-cell stages were manually transferred into a droplet of egg buffer containing 20-30 20-μm polystyrene microspheres (Polyscience) on a coverslip using an aspirator tube assembly (Sigma-Aldrich) under a dissecting microscope (Nikon SMZ745). Third, embryos on the slide were adjusted using an eyelash to arrange them in clusters of two to three embryos. Finally, the slide was covered with an 18 × 18 mm coverslip and sealed with melted Vaseline.

3D time-lapse imaging was performed under a 60X objective (PLAPON 60XO) at 20 °C ambient temperature using a spinning disk confocal microscope (Revolution XD). The microscope was equipped with an inverted microscope frame (Olympus IX73), a spinning-disk unit (Yokogawa CSU-X1) an XYZ stage with Piezo-Z positioning (ASI), an integrated solid-state laser engine (Coherent), and an EMCCD (Andor iXon Ultra 897). Images were taken using the multidimensional acquisition module of the MetaMorph software (Molecular Devices) at 300 time points, one every 75 seconds. For each time point, three slide positions were scanned for 30 Z focal planes with 1 µm spacing. Imaging parameters (laser power and exposure time) were carefully optimized to gain a reasonably high signal-to-noise ratio while minimizing photodamage to the embryos. Various parameters were tested to allow wild-type embryos to hatch on time and without obvious phenotypic abnormality relative to embryos not exposed to laser excitation. In addition, to compensate for the decay of the fluorescence signal, laser power for both mCherry and GFP was set to increase 3% for every Z plane when the focal plane extended deeper into the samples. Laser power and exposure time used for mCherry and GFP/mNeonGreen for the first Z plane were 8% for 50 *ms* and 8% for 20 *ms*, respectively.

#### Cell identification, lineage tracing, and manual curation

4D image data were organized as two-channel 3D tiff stack images for individual time points and were directly used for downstream image processing. Images were processed with the StarryNite program (Santella et al., 2014; Santella et al., 2010) for automated cell identification and tracing to reconstruct embryonic cell lineages. First, a hybrid blob-detection algorithm was used to automatically identify cells at each time point by segmenting nuclei in the 3D image stacks based on ubiquitously–expressed, nucleus-localized mCherry fluorescence (Santella et al., 2010). Second, a semi-local neighborhood-based algorithm was used to automatically trace cells over time and detect cell divisions (Santella et al., 2014).

The accuracy of cell detection and tracing is very high (> 99%); however, the accumulative nature of the errors significantly affects the accuracy of cell lineage determination at the final stage (Santella et al., 2014). Hence, to ensure high accuracy, the raw results of cell identification and tracing were subjected to extensive manual inspection and editing by two or more curators, independently, using the AceTree program (Katzman et al., 2018). Detailed procedures for lineage error detection and correction were described previously (Du et al., 2015b). Because the *C. elegans* cell lineage is invariant, most errors can be readily identified by visual or computational screening of the unusual lineage topologies. The AceTree visualizes the cell lineage results as a binary tree and provides a navigation interface to link all cells on the tree to the raw images and to correct lineage tracing results.

Finally, all traced cells were assigned a unique name following Sulston’s nomenclature (Sulston et al., 1983). Detailed nomenclature information is described elsewhere (Santella et al., 2010; Sulston et al., 1983). Briefly, cell identities were first determined for all cells (ABa, ABp, EMS, and P_2_) at the 4-cell stage based on the stereotypical cell arrangement and differential timing of cell divisions. Then, the names of their descendants were determined based on the name of the mother cell and the cell division pattern leading to the descendant cell. Cell divisions of *C. elegans* embryonic cells are classified into three types according to the orientation of cell division relative to the body axis, namely anterior-posterior (a/p), left-right (l/r), and dorsal-ventral (d/v). The full name of the mother cell is propagated to the daughter cells with an additional letter denoting the position of the daughter cell relative to the body axis following division. For instance, ABpr specifies the daughter cell of ABp that is placed on the right side following the l/r division, and ABprp specifies the daughter cell of ABpr that is located posteriorly following the a/p division. The naming of all cells follows this general rule, except for the few early blastomeres (MS, E, C, D, P_3_, P_4_, Z2, and Z3) to which special names were given in recognition of their developmental potential.

For each reporter strain, the entire cell lineage was traced until the 350-cell stage for two or more embryos (experimental replicates) with one of the embryos further traced until the 600-cell stage, when most of the terminal cells had existed for at least 15 minutes (bean stage). After lineage tracing and curation, we randomly selected 50 traced terminal cells from 50 embryos and manually examined the correctness of all cell tracking steps (n = 13,090) leading to the selected terminal cells. Only five cell tracing steps were found to be incorrect, producing an estimated tracing accuracy of 99.96%, and none of the incorrect assignments affected the expression status of corresponding TFs.

#### Quantification of reporter expression in lineaged cells

Each reporter strain used to quantify the protein expression of TF carried two fluorescent proteins. Ubiquitous mCherry was used for cell lineage reconstruction as mentioned above, and the fluorescent intensity of GFP/mNeonGreen fused to a specific TF was used to quantify the protein level of the TF in each traced nucleus. With the nuclei identified and traced following lineage construction, GFP/mNeonGreen expression in each cell can be straightforwardly determined.

GFP/mNeonGreen expression in each nucleus was measured as the average raw fluorescent intensity of all pixels within each nucleus minus the average fluorescent intensity of local background at each time point (E_n,t_ = R_n,t−_ B_n,t_). R_n,t_ was approximated by the average intensity of the fluorescence signal in the center Z plane of each nucleus at each time point. A previously described approach was used to determine B_n,t_ as the average intensity within an annular area between 1.2-radius to 2-radius from the centroid of the nucleus; nearby nuclei that overlapped with the annular area were not included in the background measurement (Murray et al., 2008). E_n,t_ values measured for a given cell were averaged to represent cellular GFP/mNeonGreen expression. Because the nuclear morphology at time points immediately before and after a cell division is usually not spherical, which would affect the accuracy of intensity measurements, E_n,t_ values at these time points were excluded from the calculation of average GFP/mNeonGreen expression in a cell.

#### Compensation for depth-dependent attenuation of fluorescence intensity

The attenuation of light with depth due to absorption and scattering of the excitation light and fluorescence is a fundamental problem associated with 3D fluorescence confocal imaging (Kervrann et al., 2004). Without adjustment, the measured fluorescence intensities of cells far from the microscope objective will be significantly weaker than those located near the objective (Figure S2). Because cells in the embryos are located at different depths, such depth-dependent attenuation of fluorescence intensity could significantly hinder making a reliable comparison of GFP/mNeonGreen expression between two individual cells.

Although the attenuation effect during 3D time-lapse imaging was partially alleviated by increasing the laser power with depth (3% per Z step), the influence of depth on fluorescence intensity was still significant in the acquired images. This effect is best illustrated by directly comparing fluorescent intensities in equivalent cells between different embryos of the same reporter strain that are oriented differently during imaging. Embryos on the slide exhibit two orientations, with either the ventral side of the embryo facing the objective (VNO) or the opposite (DNO). Thus, the fluorescence intensities of cells located near the ventral side of an embryo with a VNO orientation will be higher than the equivalent cells in embryos with a DNO orientation, and vice versa. As shown in Figure S2E, cellular TF expression was highly consistent between embryos with identical orientation (r = 0.94); however, expression levels differed dramatically between embryos with different orientations (r = 0.52), especially for cells located on the ventral or dorsal sides of the embryos (far from the center Z planes).

We developed a method to model and correct the residual attenuation effect on fluorescent intensity. We selected 11 fluorescence reporters that are ubiquitously expressed in all cells at a relatively constant level and modeled in each strain the changes in cellular fluorescent intensity as a function of the Z position of the cell (depth in the embryo). A linear model was determined to best explain the relationship between Z position and fluorescent intensity for all strains (R^2^ >0.9), and the magnitude of attenuation is highly dependent on fluorescent intensity (Figure S2B). We further modeled changes in the slope of attenuation with fluorescent intensity and revealed that the slope is highly predicted by cellular fluorescent intensity (R^2^ = 0.98, Figure S2C). Based on these results, we corrected the attenuation effect by first predicting the attenuation slope from the cellular fluorescent intensity and then using the slope and Z position of the cell to adjust the intensity. The performance of this strategy was verified by comparing the correlation coefficient of cellular GFP expression between experiment replicates with different embryonic orientations before and after the compensation. Evaluation of the performance using 46 reporter strains showed that the median correlation coefficient increased from 0.52 to 0.87, indicating the attenuation effect was significantly resolved using this method (Figure S2E). Thus, this method was applied to compensate for the attenuation of fluorescent intensities across Z planes and ensure reliable inter-cell comparisons of TF expression in the embryos.

#### Identification of TF-expressing cells

To model the distribution of background fluorescent intensity in non-expressing cells, we quantified cellular fluorescent intensity in the GFP channel in strains with the mCherry lineaging marker but not the GFP/mNeonGreen reporter. Whether or not a TF was expressed in a cell was determined by an expression cut-off of 3.2, at which the false discovery rate of being expressed was 0.01%; that is, the cellular expression level was set to 0 if the GFP/mNeonGreen intensity was lower than the cut-off.

#### Quantitative comparison of cellular TF expression to benchmark TFs curated from literature

To quantitatively evaluate the accuracy of single-cell expression, we compiled a list of 30 TFs from the literature whose expression is known to be restricted to cells from specific cell lineages or cells of specific tissue types (Table S3). These genes were used as a benchmark to evaluate the reliability of our TF expression atlas. The list includes a comprehensive accounting of TFs that are specifically expressed in the ABa (*tbx-37 and tbx-38*)(Good et al., 2004), P1 (*pal-1*)(Hunter and Kenyon, 1996), EMS (*med-2*)(Maduro et al., 2015), MS (*tbx-35* and *ceh-51*) (Broitman-Maduro et al., 2009), C (*vab-7*) (Ahringer, 1996) and multiple lineages (*tbx-8* and *tbx-9*) (Andachi, 2004), along with those specifically expressed in the neuron (*ceh-10* and *lim-4*) (Svendsen and McGhee, 1995; Zheng et al., 2005), pharynx (*pha-4*, *pha-2*, *tbx-2*, and *ceh-22*) (Kalb et al., 1998; Miyahara et al., 2004; Okkema and Fire, 1994; Raharjo et al., 2011), skin (*elt-1*, *elt-3*, *lin-26*, *nhr-23*, *nhr-25*) (Gilleard et al., 1999; Gissendanner and Sluder, 2000; Kostrouchova et al., 1998; Labouesse et al., 1996; Page et al., 1997), body muscle (*hnd-1*, *hlh-1* and *unc-120*) (Chen et al., 1994; Fukushige et al., 2006; Mathies et al., 2003) and intestine (*end-3*, *end-1*, *elt-7*, *elt-2*, *pqm-1*, *dve-1*, and *mdl-1*) (Dowen et al., 2016; Maduro et al., 2015; Reece-Hoyes et al., 2007; Yuan et al., 1998).

Although the expression of the above TFs is known to be cell-specific, the detailed single-cell dynamic expression patterns of these genes have not yet been systematically mapped. To circumvent this complexity and to allow a fine comparison of expression patterns, the cell tracks leading to all terminal cells were used as the units to generate theoretical expression patterns of the above TFs and to evaluate to what extent the cellular protein and mRNA expression data recapitulate their known patterns. The theoretical expression patterns were generated based on the known cell-specificity in terminal cells of the above TFs, with a TF being considered expressed in the cell tracks that lead to the corresponding terminal cells (Figure 2D and Table S3). The theoretical expression patterns were then used as a reference to evaluate the measured cellular protein or mRNA expression. For the testing data, a TF was considered to be expressed in a cell track if it was expressed in two consecutive cell generations within the cell track spanning from the common ancestral cell of corresponding cell lineages or tissues to the terminal cell (Figure 2D and Table S3). This cell track expression status was compared with the reference pattern to calculate the true-positive (TP), false-positive (FP), and false-negative (FN) rates. Sensitivity and precision were measured as TP/(TP+FN) and TP/(TP+FP), respectively.

#### Identification of TFs with cell-specificity

Binary cellular expression of TFs was used to identify TFs exhibiting tissue-, lineage-, and time-specificities (Table S4).

##### Tissue-specific TFs

Tissue-specific TFs were identified by assessing whether the TF is preferentially expressed in cells of a given tissue type or in cells that tend to give rise to cells of a given tissue type. Based on the cell lineage tree and tissue annotations of terminal cells, we classified all terminal cells into tissue types and calculated the probability of each cell to differentiate into each tissue type. For example, if a cell is a terminally differentiated cell of tissue-A, the probability of it being tissue-A is 1. If a cell is a progenitor cell that gives rise to four cells with one of them belonging to tissue-A, the probability of the progenitor being tissue-A is 0.25. To assess if a TF is tissue-A-specific, we compared the average tissue-A probability across all TF-expressing cells to the expected probability, which was estimated based on the number and stage (generation) of TF-expressing cells. A TF was defined as tissue-A-specific if the fold enrichment of its tissue-A probability was significantly higher than the expectation (fold enrichment > 2 and Benjamini-Hochberg corrected *p* < 0.05, hypergeometric test). Although some TFs are highly specifically expressed in a small number of cells belonging to a given tissue type, the small cell number prevented these TFs from reaching statistical significance. To accommodate this circumstance, a TF was also considered to be tissue-specific if 80% or more of the expressing cells in the terminal generation belonged to a single tissue type. In total, 101 tissue-specific TFs were identified.

##### Lineage-specific TFs

Lineage-specific TFs were identified by assessing whether the frequency of TF-expressing cells in all cells from a cell lineage is significantly higher than that expected given the size of the lineage and the total number of expressing cells (fold enrichment >2 and hypergeometric test, Benjamini-Hochberg corrected *p* < 0.05). Given that a TF could be time-specifically enriched in certain cell lineages, a TF was also considered to be a lineage-specific TF if its expression in a generation of cells from a given lineage met the above criteria. If a TF was identified to be specifically expressed in multiple inclusive cell lineages, only the largest cell lineage was kept. For example, if a TF is specifically expressed in the ABa, ABal, and ABala cell lineages, we only considered the ABa lineage. In total, 84 lineage-specific TFs were identified. It should be noted that because cells from certain lineages are biased towards specific tissue fates, a considerable number of TFs are inevitably found to be both lineage- and tissue-specific. For example, TFs that are specifically expressed in intestinal cells or their precursors should also be identified as E lineage-specific TFs. Whenever a distinction between the two was necessary, specific rules were designed and applied accordingly.

##### A-P asymmetric TFs

A-P asymmetric expression of TFs was identified by comparing the frequency of TF-expressing cells between two cell lineages following an A-P cell division. Considering that some TFs exhibit temporal specificity, the frequencies of TF-expression cells were calculated for cells at each generation and the maximum value was used to represent the expression frequency of a TF in a cell lineage. We first identified the cases in which a TF is differentially expressed between anterior and posterior cell lineages (absolute difference in expression frequency over a specific threshold). A-P asymmetric TFs were then identified if the probability of being anterior- or posterior-biased expression was significantly higher than the expectation (binomial test, *p* < 0.05). Three different thresholds of difference in expression frequency were tested (0.25, 0.5, and 0.75), and a TF was considered to be A-P asymmetric only if its expression was consistently identified to biased towards cells from one or the other lineage under two or more of the three thresholds. In addition to the lineage-based comparison, we also compared TF expression levels between individual anterior and posterior daughter cells following each cell division. A TF was considered to be A-P asymmetric if a significant difference in expression was detected between anterior and posterior cells (Wilcoxon signed-rank test, *p* < 0.01). This analysis revealed only one such TF, POP-1, the well-known regulator of asymmetric cell division. In total, 16 A-P asymmetric TFs were identified.

##### Transient expression

Transient expression of TFs was identified if TF expression was present in the mother cell but absent in one of the two daughter cells. A transient TF was identified if transient expression signals were detected in ≥ 10 mother-daughter comparisons, or the frequency of transient expression was ≥ 10%. In total, 58 TFs were identified to exhibit transient expression.

##### Quantification of developmental potential of progenitor cells

Developmental potential of each progenitor cell is defined as the distribution of tissue types across all terminal cells it generates, which was measured as information content 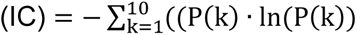, where *k* denotes each tissue category and *P*(*k*) denotes the fraction of cells within each category. Terminal cells were classified into seven categories for this calculation, comprising the five somatic tissues, cells undergoing programmed cell death, and the germ cells. The degree of fate transition between mother and daughter cells was then quantified as the reduction of IC in a daughter cell relative to the mother.

#### RNAi treatment

RNAi-mediated gene knockdown was performed following the standard feeding protocol using RNAi clones from the *C. elegans* RNAi library constructed by Dr. Julie Ahringer’s group (Source BioScience) (Kamath et al., 2001). Briefly, 10-15 worms synchronized at the L1 stage (P0) were transferred to RNAi plates containing 3 mM IPTG and seeded with bacteria expressing double-stranded RNA against a target gene of interest. After several days of RNAi exposure, embryos of P0 or P1 animals were used for imaging and phenotypic analysis. The correctness of inserts in all RNAi clones used in this study was verified by end sequencing.

#### Induction of progenitor fate transformation

We induced fate transformations between progenitor cells to determine whether the expression patterns of tissue- or lineage-specific TFs are coupled to cell fate. If coupled, we expected that the expression patterns of these TFs would change according to the new fate. Fate transformations were induced by knocking down genes known to participate in fate choice and whose loss induces the desired fate transformation (Bowerman et al., 1992; Draper et al., 1996; Du et al., 2014; Moskowitz and Rothman, 1996; Schubert et al., 2000). We performed RNAi and induced four types of progenitor cell fate transformations: (i) ABalp-to-ABarp and ABara-to-ABalp transformation by *lag-1(RNAi)*, (ii) MS-to-C transformation by *skn-1(RNAi)*, (iii) ABx-to-EMS transformation by *mex-5(RNAi)*, and (iv) ABxxx-to-C fate transformation by *mex-3(RNAi).* Four tissue-specific TFs were randomly selected for validation of the changes of their cellular expression patterns upon fate transformation, said validations consisting of ABara-to-ABala (CEH-32 and LIN-32), ABalp-to-ABarp (CEH-13), and MS-to-C transformations (TBX-7) (Figure S3D). The coupling between TF expression and tissue fate of M03D4.4 was confirmed by inducing ABx-to-EMS and ABxxx-to-C transformations (Figure S5B).

#### ChIP-seq targets of TFs

Genome-wide binding patterns and potential targets of TFs were obtained from the model organism Encyclopedia of Regulatory Networks (ModERN) database (Kudron et al., 2018). Only ChIP-seq datasets for embryo samples and with >100 target genes identified were considered.

#### Construction of TF network

We constructed a TF network by linking TFs with similar cellular expression or with a direct regulatory relationship. We first measured similarity in cellular expression between TFs using mutual information, which is widely used in probability and information theory to measure the dependence of two variables. Specifically, for each TF pair (x, y), we measured the pointwise mutual information (PMI), defined as

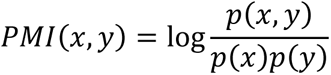

where *p(x)* is the probability of observing TF-x expression in a cell, *p*(*y*) is the probability of observing TF-y expression in a cell, and *p*(*x*, *y*) is the probability of observing co-expression of TF-x and TF-y in a cell. Two TFs are linked in the network if the observed probability of co-expression is 1.5 fold that expected and the Benjamini-Hochberg corrected *p*-value is less than 0.05 (hypergeometric test). Furthermore, we drew upon the previously generated genome-wide binding profiles of 90 TFs to link two TFs in a network if one bound the promoter of the other. Detailed information on the TF network is provided in Table S5.

#### Construction of a spatiotemporal regulatory cascade of cell lineage differentiation

The spatiotemporal regulatory cascade of TFs was constructed in three steps: (i) dividing cell lineage differentiation into spatiotemporal regulatory modules, (ii) assigning TFs to corresponding regulatory models, and (iii) extracting regulatory relationships between TFs from the TF network. The cascade is provided in Table S5.

##### Deconstruction of cell lineage differentiation into spatiotemporal modules

We deconstructed the regulation of cell differentiation as the regulation of differentiation of each tissue type in tissue progenitor cells from different cell lineages (spatial modules). To do so, we classified all terminally differentiated cells into five major tissue types: the neuronal system, pharynx, skin, body muscle, and intestine. We then performed clonal analysis as done previously to identify all tissue progenitor cell lineages that differentiate exclusively into a given tissue type (Du et al., 2014). The clonal analysis was performed bottom-up based on the tissue types of all terminal cells to quantify the probability of each progenitor cell differentiating into a specific tissue type. For each progenitor cell, this probability was calculated recursively by averaging the score of its two daughter cells. The highest-scoring clone with a high probability (≥ 0.85) of producing a considerable number of terminal cells (three or more) that belong to the same tissue type was defined as a tissue progenitor cell lineage. The clonal analysis identified ABala cell (one of the cells at the 12-cell stage embryo) as the founder cell of a neuronal cell lineage as most of terminal cells from this lineage differentiates into neurons or glia cells. However, considering that it is highly unlikely that neuronal fate specification occurs as early as the 12-cell stage, we manually divide this large lineage into four smaller ones (ABala-aa, ABala-ap, ABala-pa, and ABala-pp). In total, we identified 56 tissue progenitor cell lineages: 24 for the neuronal system, 11 for the pharynx, 13 for the skin, seven for body muscle, and one for intestine.

Each tissue progenitor cell lineage was then divided into three temporal modules based on the maturity of cell differentiation. First, development from the zygote to the mothers of the founder cells of identified tissue progenitor cell lineages was considered to comprise upstream regulation of tissue differentiation (temporal module I). Second, development from the founder cell to the mothers of terminal cells is considered to comprise the specification/early differentiation of tissue fate (temporal module II). Finally, generation of the terminal cell is considered the terminal differentiation of tissue fate (temporal module III). If a progenitor cell lineage contained three or fewer generations of cells, only two temporal modules were considered, specification/early and terminal differentiation.

##### Assignment of TFs into spatiotemporal modules

All tissue-specific TFs, including those also identified as lineage-specific ones, were assigned to temporal module II or module III of the corresponding tissue progenitor cell lineages based on their expression time. If a tissue-specific TF was expressed only in the terminally differentiated cells of corresponding tissues, it was assigned to temporal module III. If a tissue-specific gene was expressed in cells from the founder of each tissue progenitor cell lineage to one generation earlier than the terminally differentiated cell, that TF was assigned to temporal module II. All exclusive lineage-specific TFs that were expressed in cell generations no later than the daughter cells of the founder cell of each tissue progenitor cell lineage were assigned to module I.

##### Extracting regulatory relationships between TFs from the TF network

Finally, TF relationships were extracted from the TF network to generate subnetworks revealing potential regulatory relationships between TFs within each spatiotemporal module and between different modules of the same tissue type.

#### Comparison of cellular regulatory states (CRSs)

Divergence of cellular regulatory state (CRS) between cells was quantified as the Jensen-Shannon divergence of TF expression. Ubiquitously expressed TFs were excluded from the calculation.

#### Gene enrichment analysis

Enrichment of Gene Ontology terms and RNAi phenotypes in TF target genes were performed using the Gene Set Enrichment Analysis tool of WormBase (https://wormbase.org/tools/enrichment/tea/tea.cgi) with default parameters (Angeles-Albores et al., 2018).

#### Single-cell phenotypic analysis

##### Cell lineage and TF expression changes

Cell lineage tracing and TF expression quantification in mutant embryos were performed as in constructing the TF expression atlas. Programmed cell death was determined by a characteristic sequence of morphological changes in the nucleus of the corresponding cell and the disappearance of the nucleus.

##### Cell position changes

The definition and quantification of relative cell positions were performed as previously (Li et al., 2019). Briefly, a vector of the geometric distance between a target cell and all other co-existing cells at the same developmental stage (reference cells) was used to define the relative position of the target cell. Relative cell positions were then compared between equivalent cells in perturbed and wild-type embryos and in individual wild-type embryos to determine whether a cell in the perturbed embryo deviated significantly from the normal distribution of the corresponding wild-type cells. The similarity of relative cell positions was calculated as the root-mean-square deviation (RMSD) score of the distance vectors. Twenty-eight previously processed wild-type embryos were used to calculate the Z-scores of relative cell positions in the perturbed embryos (Li et al., 2019), and a cell with a Z score over 2.576 (*p* < 0.01, two-tailed) was considered as having changed cell position.

## SUPPLEMENTARY FIGURES LEGENDS

**Figure S1.**
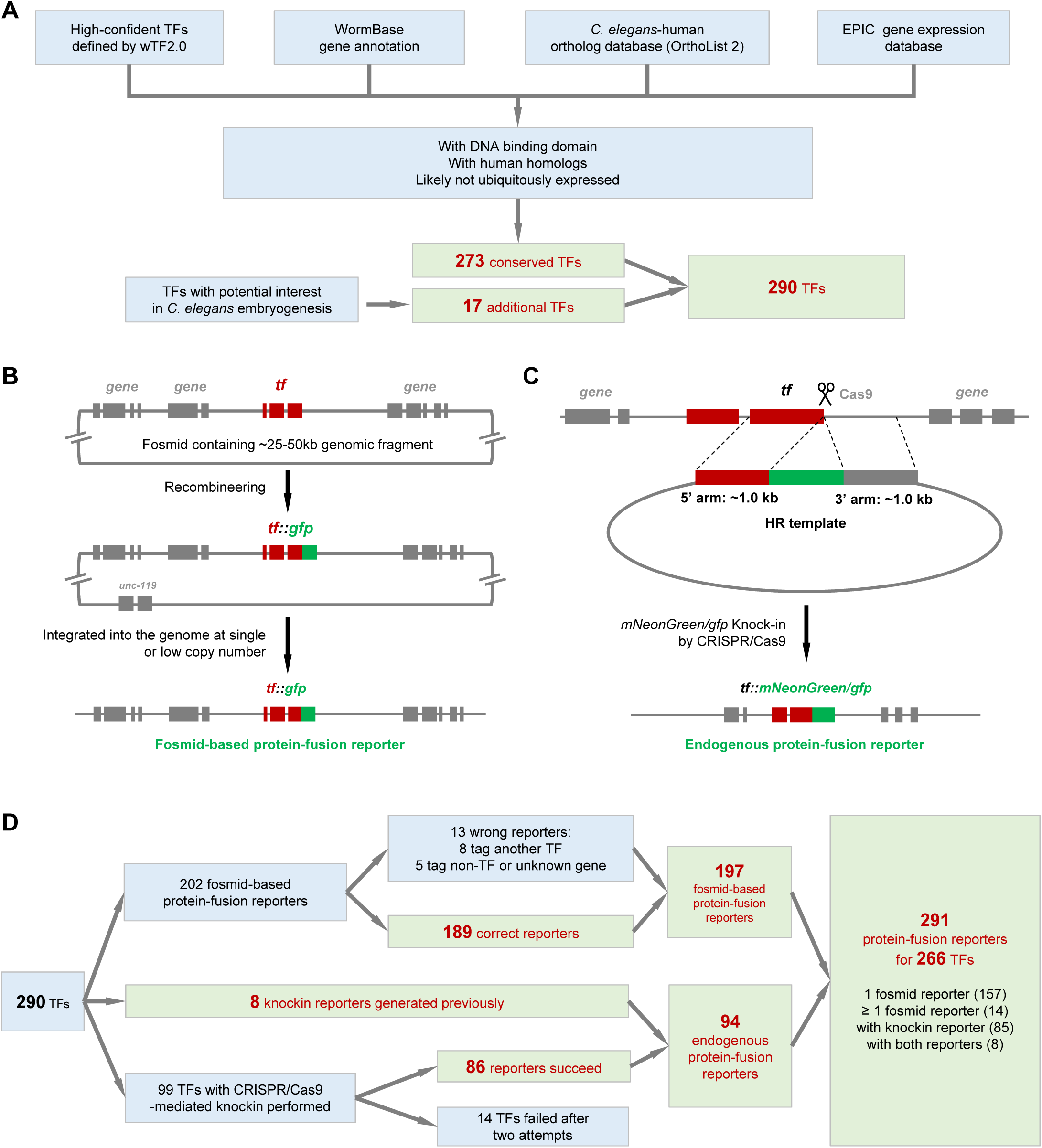
Generation of protein-fusion fluorescent reporter of TFs, Related to Figure 1. (A) Selection of 290 high-confidence TFs as target genes, including 273 TFs that have human homologs and 17 worm-specific TFs with potential interest in cell differentiation. (B) Fosmid-based protein-fusion reporters. 202 protein-fusion reporters of 185 TFs have been generated by the ModENCODE projects in which GFP (green) was fused to a TF (orange) in a fosmid by recombineering, and the fosmid was then integrated into the *C. elegans* genome (Araya et al., 2014; Sarov et al., 2012). (C) Tagging endogenous TFs with mNeonGreen/GFP by CRISPR/Cas9-mediated gene knockin. A homologous recombination (HR) template sequence containing the coding sequence of mNeonGreen/GFP (green) flanked by 800-1000 bp sequence homologous to the endogenous TF locus was used to guide HR following cleavage by Cas9 at the N- or C terminals of the TF gene. (D) Flowchart showing the sources of 291 protein-fusion reporters for 290 TFs.

**Figure S2.**
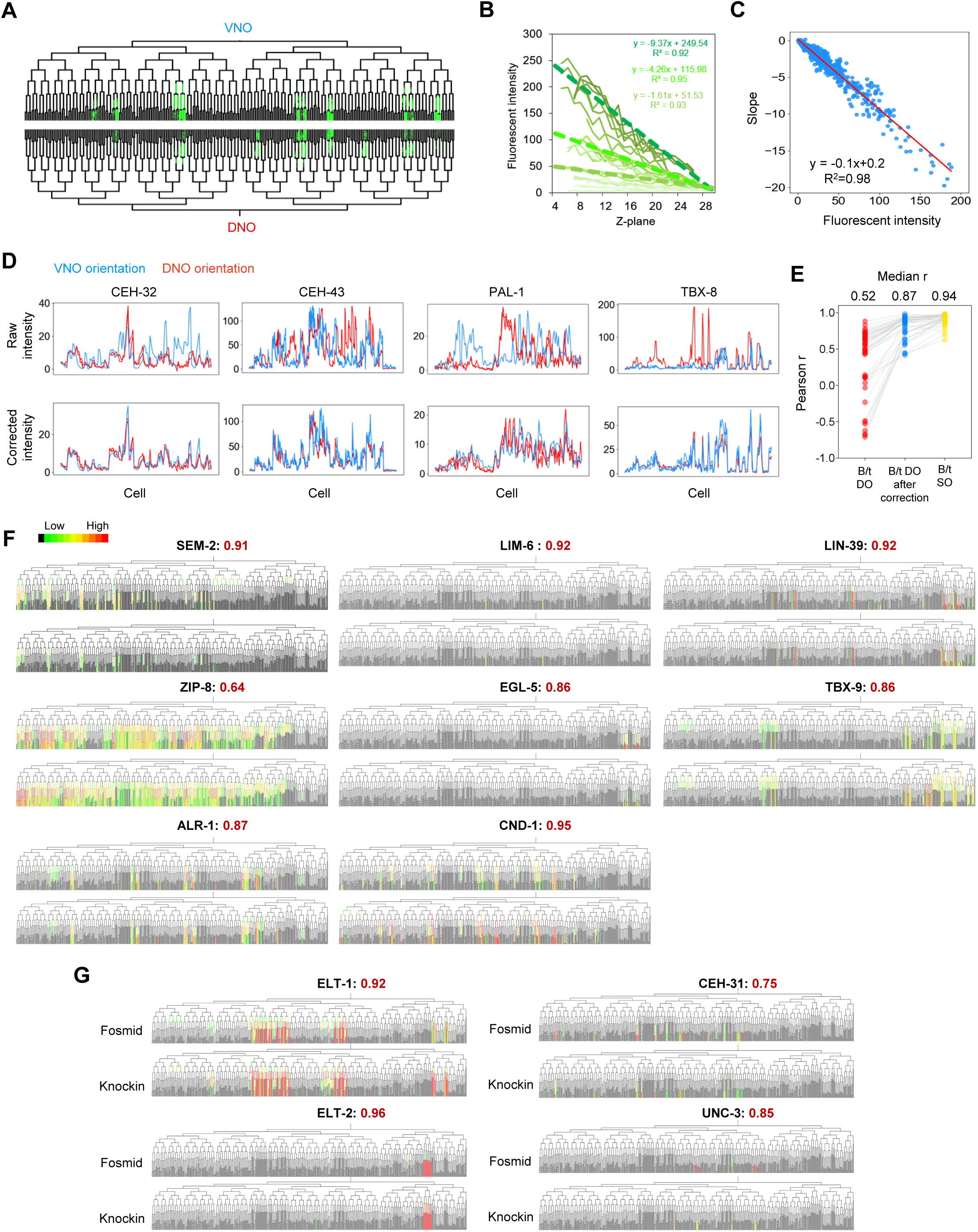
Quality assessment of cellular quantification of TF expression, Related to Figure 2. (A) Tree visualization of cellular TF expression levels (green) in two embryos of the same strain but oriented differently (VNO and DNO) during imaging. (B) Decrease of mNeonGreen/GFP fluorescent intensities following the increase of cell depth in the sample. TF reporter strains showing ubiquitous and relatively homogenous cellular expression were used to model the relationship between sample depth (Z plane) and attenuation of fluorescent intensity. Dashed lines indicate the results of linear regression for TF strains exhibiting different average expression levels. (C) Dependency of the attenuation effect on the level of fluorescent intensity. Red line indicates the result of linear regression. (D) Representative examples showing the performance of fluorescent intensity correction across depth. Top and bottom panels show TF expression in equivalent cells of two embryos with different orientations, before and after correcting the attenuation effect. (E) Comparison of the Pearson correlation coefficient of TF expression in equivalent cells between different embryos. ‘B/t DO’ and ‘B/t DO after correction’ denote the comparison between embryos with different orientations without and with correction, respectively. ‘B/t SO’ indicates the comparison between embryos with the same orientation. (F) Comparison of cellular TF expression between different fosmid-based protein-fusion reporter strains that tag the same TF. (G) Comparison of cellular TF expression between fosmid-based (top) and knockin-based (bottom) reporter strains that tag the same TF.

**Figure S3.**
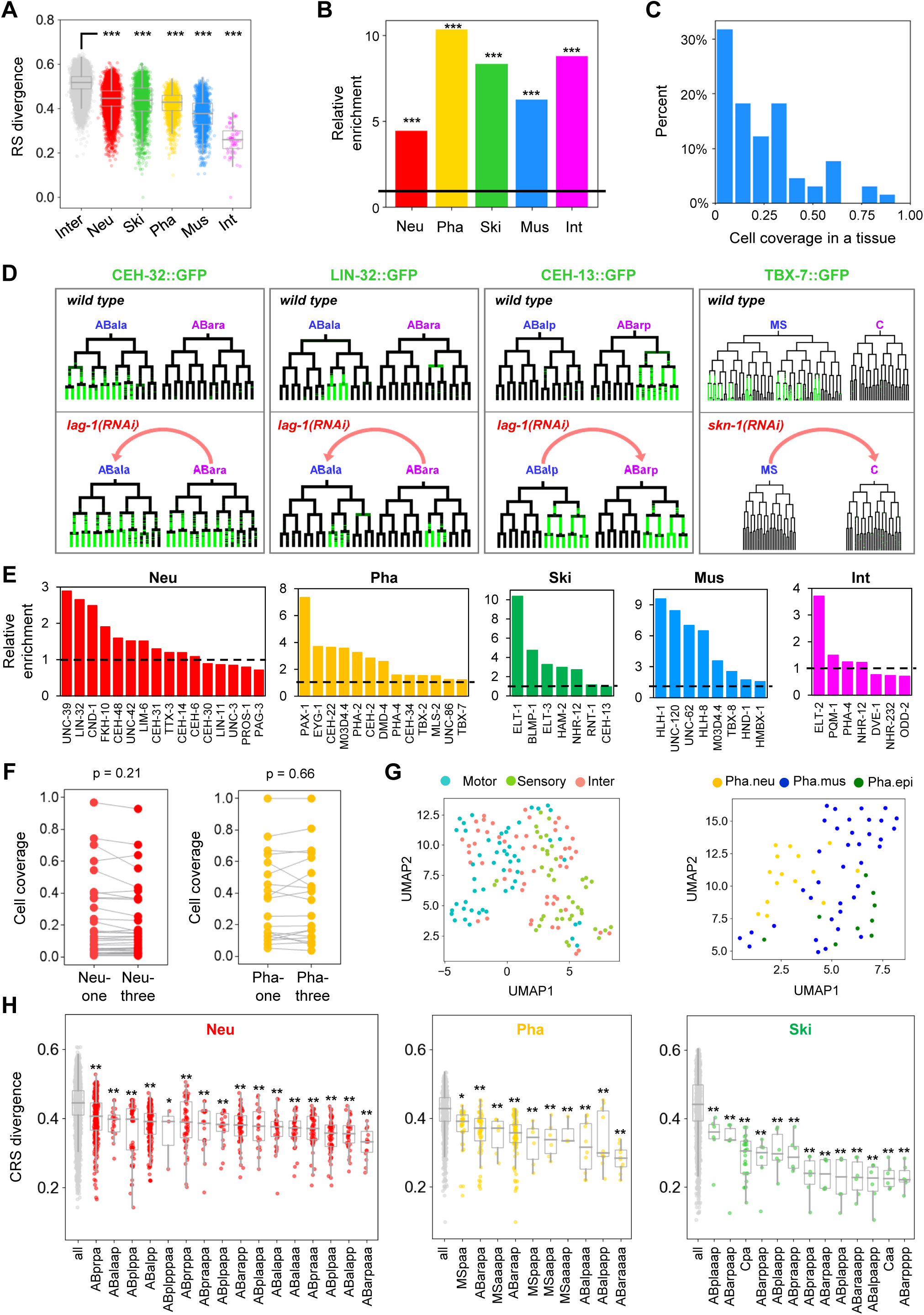
Lineage-restricted expression of tissue-specific TFs, Related to Figure 3. (A) Inter- and intra-tissue divergences of CRSs between cells. Statistics: Mann–Whitney U test. *** indicates *p* < 0.001. (B) Enrichment of the overlap between tissue-specific TFs identified in this study and by (Warner et al., 2019). Statistics: Hypergeometric test, *** indicates *p* < 0.001. (C) Distribution of cell coverage of tissue-specific TFs in corresponding tissue types. (D) Tree visualization of the cellular expression patterns of TFs before (top) and after (bottom) switching progenitor cell fate by RNAi against specific fate specifiers. Arrows indicate fate transformations. (E) Relative enrichment of tissue-specific genes in the targets of tissue-specific TFs. In each tissue, relative enrichment was measured as the ratio of the frequency of genes specifically expressed in this tissue to that of genes specifically expressed in other tissues. Tissue-specific genes identified by (Warner et al., 2019) were used. (F) Cell coverage of neuronal- and pharyngeal-specific TFs in corresponding tissues before and after dividing the tissue cells into subtypes. Statistics: Wilcoxon signed-rank test. (G) UMAP plot of neuronal (left) and pharyngeal (right) subtypes based on TF expression. Parameters: min_dist = 0.8, n_neighbors = 5. (H) Comparison of intra-tissue CRS divergences between all cells and between cells of each tissue progenitor cell lineage. Statistics: Mann–Whitney U test. ** and * indicate *p* < 0.01 and *p* < 0.05 respectively.

**Figure S4.**
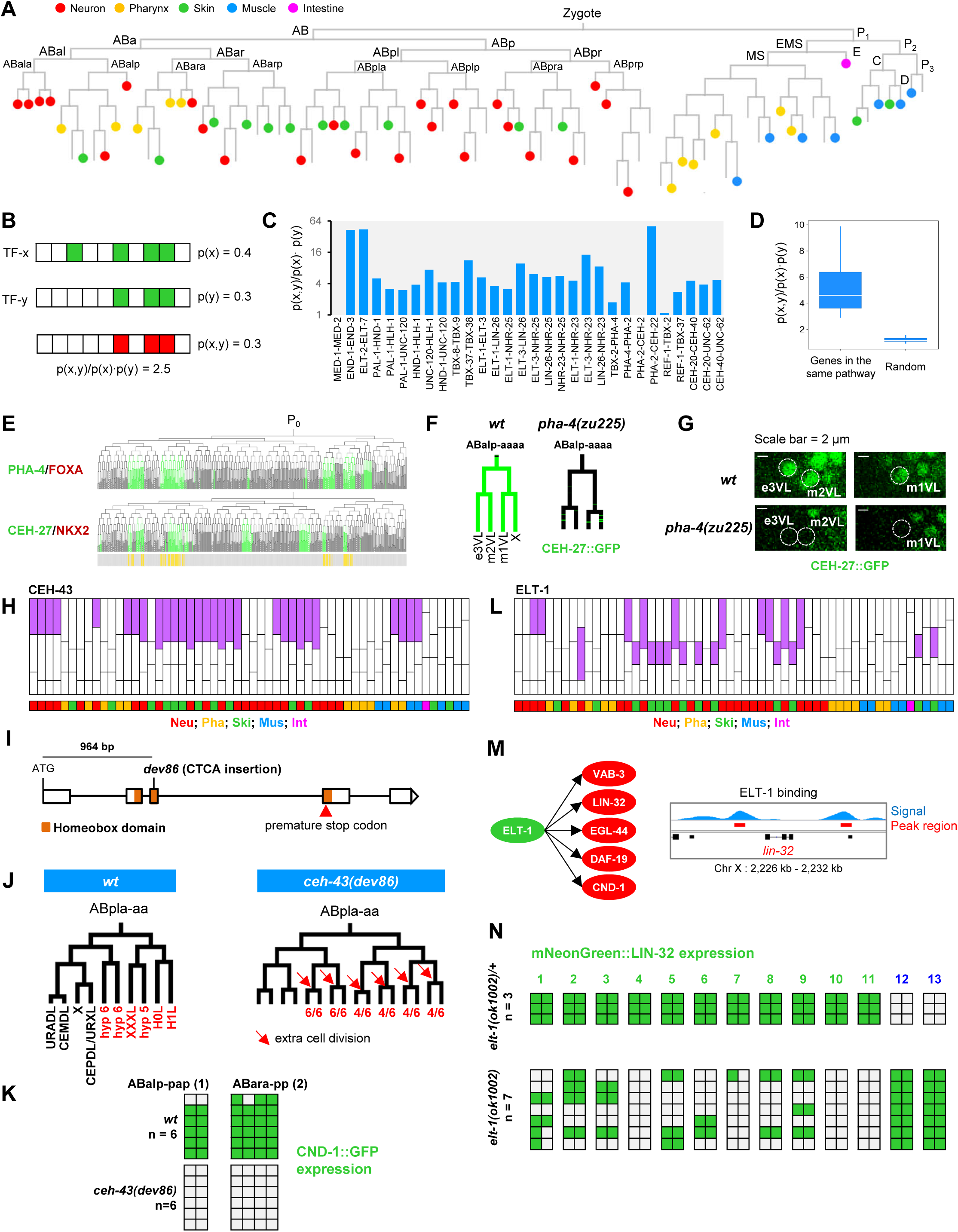
Construction of a spatiotemporal TF cascade reveals functions of CEH-43 and ELT-1 in lineage differentiation of neuronal cells, Related to Figure 5. (A) Tree visualization of the cell lineage leading to all progenitors of defined tissue types. (B) An example of calculating the similarity in single-cell expression of a given gene. (C) Single-cell expression similarity between 30 pairs of TFs known to function in the same pathway(Good et al., 2004; Maduro et al., 2015; Morck et al., 2004; Neves and Priess, 2005; Smith and Mango, 2007; Van Auken et al., 2002; Yanai et al., 2008). (D) Comparison of expression similarity between genes in the same pathway (n = 30) and between randomly selected gene pairs (n = 30). Statistics: Mann–Whitney U test. (E) Single-cell protein expression of PHA-4 and CEH-27 in lineaged cells. (F) Comparison of CEH-27 protein expression in the ABalpaaaa lineage between *wt* and *pha-4(zu225)* embryos. (G) Micrographs showing the loss of CEH-27 expression in *pha-4(zu225)*. (H) The inferred spatiotemporal function of CEH-43. (I) Generation of a *ceh-43* loss of function allele. A 4-bp insertion (CTAG) that introduces a premature stop codon into the open reading frame was induced 964 bp downstream of the start codon of *ceh-43* by CRISPR/Cas9-mediated gene editing. (J) Comparison of cell division pattern of the ABplaaa lineage in the *wt* and *ceh-43(dev86)* embryos. (K) Comparison of single-cell CND-1 protein expression in corresponding cell lineages of individual *wt* and *ceh-43(dev86)* embryos. (L) The inferred spatiotemporal function of ELT-1. (M) Left: All neuronal-specific TFs targeted by ELT-1. Right: ELT-1 binding pattern near the gene *lin-32*. (N) Comparison of single-cell LIN-32 protein expression in individual *elt-1(ok1002)/+* and *elt-1(ok1002)* embryos in corresponding cell lineages.

**Figure S5.**
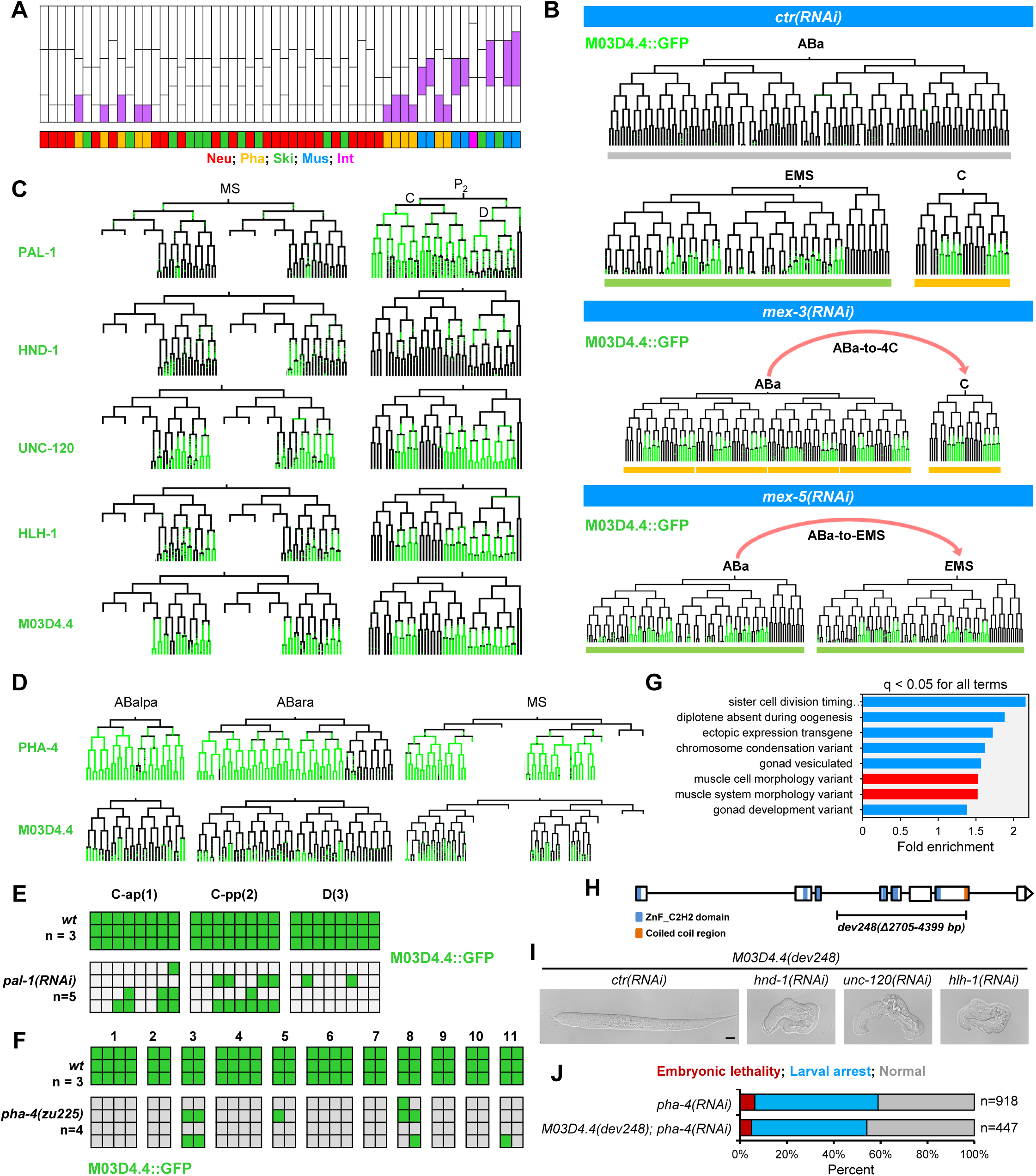
M03D4.4 participates in muscle differentiation in the body wall and pharynx, Related to Figure 6. (A) The inferred spatiotemporal function of M03D4.4. (B) Single-cell protein expression of M03D4.4 in cell lineages before and after switching fates of specific progenitor cells. The homeotic transformation of ABala, ABalp, ABara, and ABarp cell fates to C (ABa- to-4C) was induced by *mex-3(RNAi)*. The homeotic transformation of ABa cell fate to EMS (ABa-to-EMS) was induced by *mex-5(RNAi)*. (C) Comparison of single-cell expression of M03D4.4 and other known regulatory TFs (PAL-1, HND-1, UNC-120, HLH-1) in the cell lineages that produce body muscle (MS, C, and D). (D) Comparison of single-cell protein expression of M03D4.4 and PHA-1 in three cell lineages (ABalpa, ABara, and MS) that produce pharyngeal muscle. (E) Comparison of single-cell M03D4.4 protein expression in corresponding cell lineages between individual *wt* and *pal-1(RNAi)* embryos. (F) Comparison of single-cell M03D4.4 protein expression in corresponding cell lineages between individual *wt* and *pha-4(zu225)* embryos. (G) Enrichment of RNAi phenotypes in M03D4.4 target genes. Statistics: Hypergeometric test, Benjamini-Hochberg corrected *p*-value. (H) Generation of a *M03D4.4* loss of function allele. A 1695-bp deletion that removes exons 4-6 and most part of exon 7 was induced by CRISPR/Cas9-mediated gene editing. (I) Representative differential interference contrast (DIC) images showing synthetic phenotypes between *M03D4.4* and three genes regulating body muscle differentiation. (J) Comparison of embryonic lethality and larval arrest phenotypes between *pha-4(RNAi)* (n = 918) and *M03D4.4(dev248); pha-4(RNAi)* (n = 447) animals.

**Figure S6.**
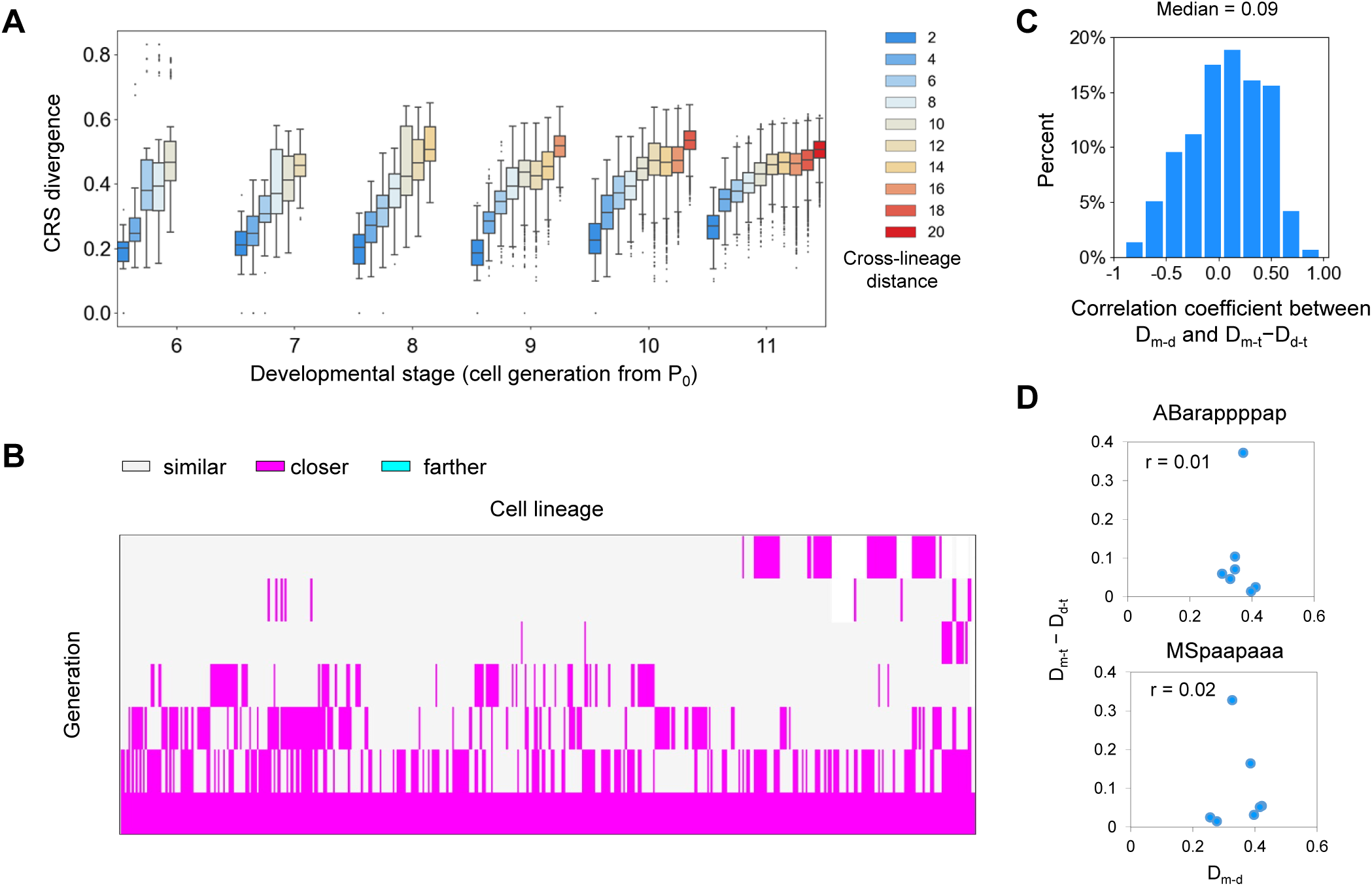
CRS dynamics during cell differentiation, related to Figure 7. (A) Change of CRS divergence as a function of cross-lineage distance between cells at different developmental stages. (B) Classification of cell divisions into three categories according to whether or not a mother-to-daughter CRS transition drives CRS toward the terminal CRS. If CRS divergence between a mother and daughter cell (D_m-d_) was larger than the divergence between the initial and terminal cells in the cell track divided by the total number of cell divisions (D_i-t_/Nd), the cell division was classified as either closer or father according to its direction. Otherwise, the cell division was classified as not driving CRS either towards or away from the terminal CRS (similar). (C) Distribution of the Pearson correlation coefficient between D_m-d_ and D_m-t_ minus D_d-t_ across all cell divisions. (D) Scatter plot showing the poor correlation between D_m-d_ and D_m-t_ minus D_d-t_ in two representative cell tracks.

**Figure S7.**
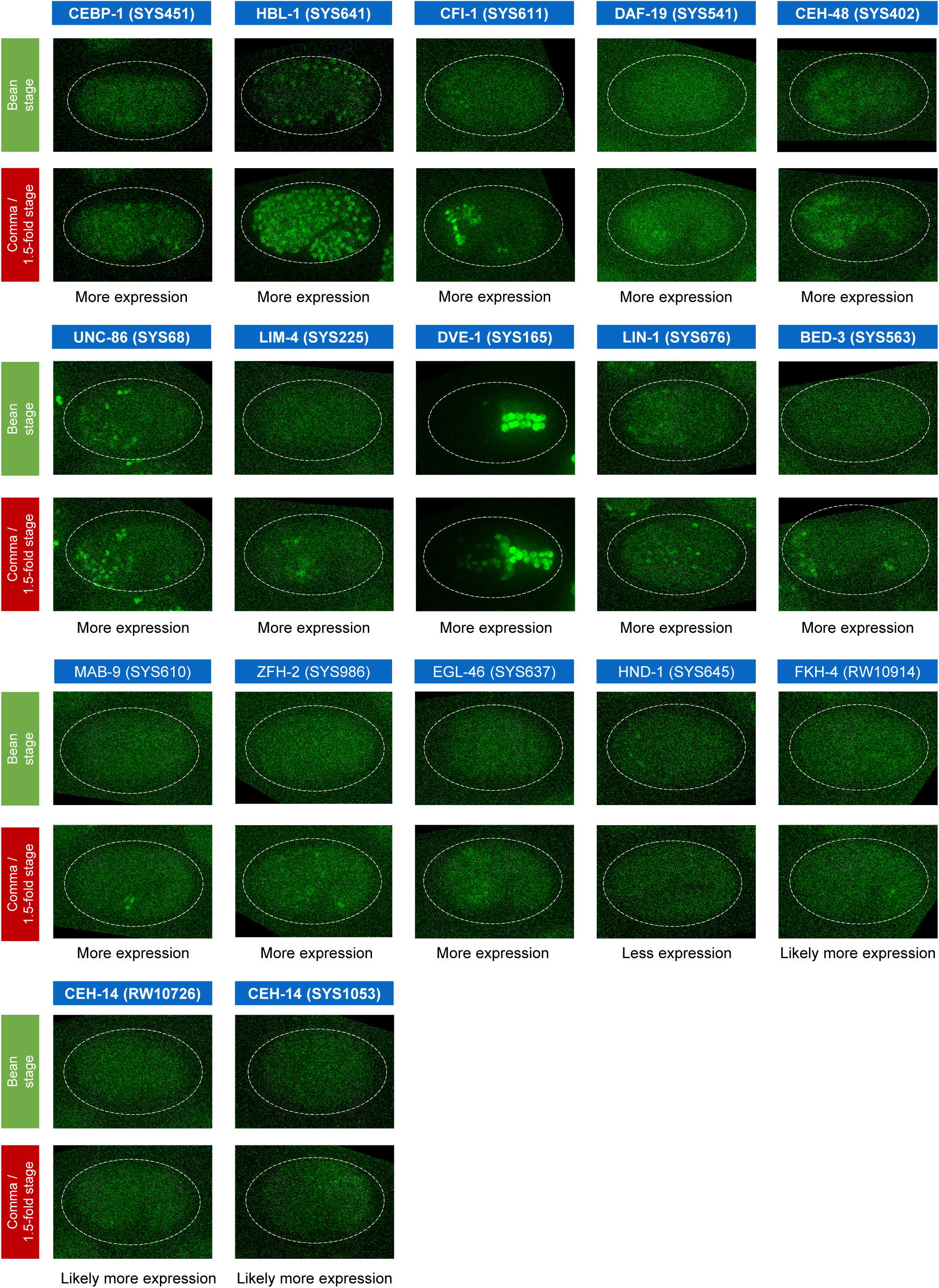
TFs potentially changed expression pattern from bean to comma/1.5-fold stage, related to Discussion Maximum projections of 3D images showing expression of 16 TFs at the end time point of cell lineage tracing (∼bean stage) and at comma/1.5 fold stage. These TFs were identified by manual inspection of 3D images.

## SUPPLEMENTARY TABLES

Table S1. Generation of dual-fluorescent reporters of TFs, Related to Figure 1.

1^st^ sheet lists the 290 selected TFs and related information.

2^nd^ sheet lists the sources and verification of fluorescent reporters used in this study.

3^rd^ sheet lists the strain names and associated genotypes of all dual-fluorescent reporters of TFs. 4^th^ sheet lists the other strains and reagents used in this study.

Tables S2. Cellular expression pattern of TFs, related to Figure 2.

1^st^ sheet lists the quantitative expression of 291 protein-fusion TF reporters (for 266 TFs) in 1,204 lineaged cells.

2^nd^ sheet lists the quantitative expression of all TFs in multiple embryos (≥ 2) for the same reporter strain up to the 350-cell stage.

Table S3. Comparison of cellular TF expression to the literature, Related to Figure 2.

1^st^ sheet lists previous RNA-seq data for the TFs identified as exhibiting sporadic expression in this study.

2^nd^ sheet lists the results of comparing the expression patterns of TFs reported in this study to those reported in the literature. We only considered TFs for which expression patterns in specific lineages, tissue types, cells, and developmental stages were previously described.

3^rd^ sheet lists the correlation coefficient of single-cell protein expression of TFs between this and previous studies.

4^th^ sheet lists the results of cell-by-cell expression comparisons for 30 TFs with well-documented cellular expression patterns.

Table S4. TFs with cell specificity, Related to Figure 3 and Figure 4.

1^st^ sheet lists all identified lineage-specific TFs, enriched tissue types, and enrichment scores.

2^nd^ sheet list the enriched RNAi phenotypes for the ChIP-seq target genes of each tissue-specific TF (n = 48). Yellow highlights those phenotypes that are related to the function of corresponding tissue.

3^rd^ sheet lists all identified lineage-specific TFs, enriched lineages, and enrichment scores.

4^th^ sheet lists all identified A-P asymmetric TFs and related bias scores.

5^th^ sheet lists TFs with transient expression. ‘1’ denotes a TF that is expressed in a mother cell but absent in at least one of its daughters.

Table S5. The spatiotemporal cascade of TFs, Related to Figure 5.

1^st^ sheet lists the assignment of TFs into spatiotemporal modules.

2^nd^ sheet lists all relationships between TFs inferred based on expression similarity and previous ChIP-seq experiments.

3^rd^ sheet lists the inferred TF cascade for each tissue progenitor cell lineage.

Table S6. Cell position and expression change phenotypes, related to Figure 5 and Figure 6.

1^st^ sheet lists the identified cell position defects in individual *ceh-43(dev86)* embryos.

2^nd^ sheet lists cellular expression levels of CND-1 in individual wild-type and *ceh-43(dev86)* embryos.

3^rd^ sheet lists cellular expression levels of LIN-32 in individual *elt-1(ok1002)/+* and *elt-1(ok1002)* embryos.

4^th^ sheet lists changes in cellular expression levels of M03D4.4 in individual embryos upon *pal-1* and *pha-4* perturbations.

